# Landscape-wide metabarcoding of the invasive bumblebee *(Bombus terrestris)* shows interactions among the gut microbiome and pollenbiome

**DOI:** 10.1101/2024.09.08.611921

**Authors:** Sabrina Haque, Hasinika KAH Gamage, Cecilia Kardum Hjort, Fleur Ponton, Francisco Encinas-Viso, Ian T Paulsen, Rachael Y Dudaniec

## Abstract

Many species of social insects introduced to regions beyond their native ranges have become highly invasive. The introduction of the eusocial European buff-tailed bumblebee, *Bombus terrestris*, to the island of Tasmania (Australia) ∼30 years ago is of concern due to its ecological impacts and its potential to spill over pathogens to native bees or commercially important honeybees. The health of *B. terrestris* is intricately connected with its gut microbiome and diet; however, environmental variables may also interact, particularly during invasion into novel environments. Using landscape-wide sampling and a metabarcoding approach to characterize the gut bacteria (16S rRNA) and diet composition from foraged pollen (ITS2: floristic diversity of pollen baskets), this study investigates how the gut microbiota of *B. terrestris* workers is affected by nutritional diversity (‘pollenbiome’) and environmental variation across diverse landscapes of its invasive range in Tasmania. Gut bacterial community composition and diversity were significantly predicted by site annual precipitation and percentage of pasture. Further, a positive interaction between site annual precipitation and site annual temperature significantly predicted gut bacterial diversity. The interaction effect of pollen diversity and average summer wind velocity was also significantly and positively related to gut bacterial diversity. Following comparison of Akaike information criterion (AIC) and sum of weights, the percentage of pasture was identified as the most strongly weighted variable, which, along with pollen diversity, had a negative impact on gut bacterial diversity. These insights help to uncover how environmental interactions affect the gut microbiome of *B. terrestris* in an invaded landscape with novel nutritional resources. This knowledge contributes to understanding the factors that predict the spread and persistence of invasive bumblebees.

## Introduction

The dynamic relationship between the gut microbiome and diet can be a key determinant of animal health in natural ecosystems (Douglas, 2018), yet the impact of the environment on this relationship is not clearly understood (e.g., Baniel et al. 2021; Loo et al. 2019). The potential of this relationship to aid in the success of invasive species is also unknown (Fontaine et al. 2020; Zhu et al. 2021). Novel diets can alter gut microbiomes of invasive species which may potentially impact their health and capacity to invade (Escalas et al., 2022). Therefore, unravelling the links between environment, nutrition, and the gut microbiome of invaders as they establish within novel locations is relevant for understanding successful biological invasions. Among terrestrial invaders, insects, especially social and eusocial Hymenoptera (e.g. bees, wasps, ants) are invasive on a global scale (Russo 2016; Manfredini et al., 2019; Ghisbain et al., 2021). There has been increasing concern about the environmental or ecological impacts of deliberate or accidental introductions of pollinators (i.e., bees) into agricultural settings beyond their native ranges (Russo, 2016; Aizen et al., 2020). Hence, elucidating how environmental factors and novel diets influence the gut microbiome of invasive pollinators is important for understanding how these invaders maintain health and persist within novel ecosystems.

Introduced pollinators generally exhibit a preference for foraging on introduced plants and they frequently serve as primary pollinators for numerous weeds (Goulson, 2003; Hanley & Goulson, 2003; Lowenstein et al., 2019; O’Connell et al., 2021). Generalist pollinators, known for their adaptability and ability to exploit a variety of resources, may be more prevalent in urban areas because of the diverse floral resources available (Zanette et al. 2005; Biesmeijer et al. 2006; Jędrzejewska-Szmek and Zych 2013). Studies suggest that urban landscapes may offer valuable opportunities for introduced pollinators as they often have a remarkable variety of floral resources, including crops in community gardens, ornamental plants and self-seeding weeds (Frankie et al. 2005; Matteson & Langellotto 2009; Hülsmann et al. 2015; Lowenstein & Minor 2016). Furthermore, annual plants may offer less reliable resources compared to perennials, and while ornamental plants may appear less attractive to pollinators (Garbuzov et al., 2015; Garbuzov et al., 2017), with weeds being potentially more appealing for foraging purposes. Interactions between plants and invasive pollinators may therefore reveal how shifts in foraging behaviour modify ecosystems and influence pollinator invasiveness.

The simple yet distinct microbial community found in eusocial bees has been associated with various health benefits (Martinson et al., 2011; Kwong & Moran, 2016). Research indicates that a diverse gut microbiota contributes significantly to aspects such as efficient digestion of plant-based foods, weight regulation through nutrient availability, physiological processes, endocrine signalling, resilience to parasites and pesticides, behaviours, and immune responses of the host (Anderson et al., 2011; Koch & Schmid-Hempel, 2011; Vásquez et al., 2012; Engel & Moran, 2013; Schwarz et al., 2016; Ricigliano et al., 2017; Zheng et al., 2017; Dosch et al., 2021). Social behaviour strongly affects the mode of acquisition of gut microbes in bees and other animals. In highly social corbiculate (pollen basket bearing) bee species, the gut microbiome consists of a relatively small and consistent group of coevolved taxa (Kwong & Moran, 2016). These microbes play key roles in digestion, growth, immunity, and detoxification (Kwong et al., 2017). In contrast, solitary bees generally host a more diverse array of gut microbes, often acquired from the environment rather than through social interactions within the nest (McFrederick et al., 2017). Some solitary bees do harbor a core group of bacteria, which can overlap with those found in social bees (Gu et al., 2023). However, the specific functions of gut microbiome in solitary bees are less understood, although microbes associated with pollen provisions are known to be important for larval nutrition and development (Vanette et al., 2012; Stefan et al., 2019; Stefan & Dharampal, 2019; Dharampal et al., 2019).

Bee health is tightly linked with their gut microbiome and diet (pollen and nectar) (Motta & Moran, 2024), however, environmental variables may also interact with bee health outcomes, which is likely to be significant when invading diverse novel landscapes (Anderson et al., 2011; Engel et al, 2016). Introduced bees may compete with native pollinators for floral resources and nesting habitats, introduce pathogens to native organisms, as well as facilitate the spread of exotic weeds which disrupts native plant communities (Goulson,2003; Abrol, 2012). While bees feed on nectar for carbohydrates, pollen serves as the main source of proteins, amino acids, lipids, starch, sterols, vitamins, and minerals, which are vital for larval rearing, physiological development, immunity, and longevity in bees (Roulston & Buchmann, 2000; Brodschneider & Crailsheim, 2010; Alaux et al., 2010; Nicolson, 2011). Diverse diets that include pollen from numerous plant species are advantageous for bee health as they offer a wider array of nutrients, compared to monofloral diets solely from one plant species (Brodschneider et al., 2021). At the colony level, pollen facilitates the production of royal jelly by young workers, which is later used for nourishing larvae, queens, drones, and older workers (Crailshem et al., 1992; Crailshem, 1992; Di Pasquale et al., 2013). In addition, pollen can influence the capacity of bees to metabolize toxic compounds, such as pesticides (Brascou et al., 2021). Genomic and metagenomic analyses have suggested that the gut bacteria of bees aid in digesting macromolecules, providing nutrients, neutralizing dietary toxins, and defending against parasites (Engel et al., 2012; Kwong et al., 2014; Lee et al., 2015, Engel et al., 2016). Hence, exploring the relationship between pollen-derived nutrition and the composition of bee gut bacterial communities may be beneficial for understanding bee health and factors that facilitate their invasion.

The use of DNA metabarcoding to characterize pollen diversity offers a powerful method for investigating the pollen composition of bee diets, and spatial and temporal fluctuations in plant-pollinator interactions (Hornick et al., 2021, Bell et al., 2022; Milla et al., 2022; Encinas-Viso et al., 2022). Pollen DNA metabarcoding can identify the multiple plant taxa visited from the pollen grains carried within bee pollen packets or attached to the body (Keller et al., 2015; Pornon et al., 2016; Bell et al., 2017). Metabarcoding studies of pollen have revealed new interactions within flower-visitor networks, uncovering missing links in the pollination biology of cryptic plant species (Pornon et al., 2017; Lucas et al., 2018; Arstingstall et al., 2021; Encinas-Viso et al., 2022). Therefore, pollen metabarcoding can enhance our understanding of the ecological and evolutionary processes that influence connections between plant and pollinator gut bacterial communities across diverse environments.

Tasmania, the island state of Australia, witnessed a remarkably swift and successful introduction of the European buff-tailed bumblebee, *Bombus terrestris*, in 1992 (Semmens et al., 1993; Hingston 2006a). The initial colonization event was believed to have involved only a small number (perhaps as low as three) of *B. terrestris* queens from New Zealand (Schmid-Hempel et al., 2007). Within a decade of its introduction into the city of Hobart, it rapidly spread throughout the entire island (Hingston et al., 2002). Despite significant isolation-by-distance and spatial variability in effective migration rates, the Tasmanian *B. terrestris* population displays high gene flow alongside low genetic diversity (Kardum Hjort et al., 2023a). *B. terrestris* workers also exhibit some morphological divergence in relation to environmental conditions across Tasmania (Kardum Hjort et al., 2023b). The widespread invasion of *B. terrestris* is believed to have adversely affected the Tasmanian ecosystem by competing with and displacing native bees and other pollinators (Hingston & McQuillan, 1999, Hingston & Wotherspoon, 2017, Debnam et al., 2021). Consequently, the invasion has led to a decline in the pollination efficiency of native plants due to nectar robbing or physical damage to flowers (Hingston & McQuillan, 1998).

Research findings indicate that invasive bumblebees can reduce available resources for managed honeybees by robbing them of raspberry flower buds, thus revealing their potential impact on honey production (Sáez et al., 2017). Furthermore, *B. terrestris* pollination is thought to facilitate the spread of exotic weeds in Tasmania (Hingston, 2006b). Although mainland Australia is currently free of *B. terrestris*, there is a risk of it spreading to the mainland due to its high mobility, the presence of suitable habitats, and climates similar to that of Tasmania, particularly within the south and east coastal regions (Hingston 2007; Acosta et al., 2016; Fijen 2021; Kardum Hjort et al., 2023a; Kardum Hjort et al., 2023b). Therefore, the introduction of *B. terrestris* to Tasmania is of concern due to both its ecological impacts and its potential to transmit pathogens to native bees and insects, and its potential to harbor pathogens from commercial honeybees.

Here we explore the intricate relationships between gut microbiome and pollenbiome diversity in the invasive bumblebee, *B. terrestris*, across varying environmental conditions in Tasmania, Australia. Through landscape-wide sampling and a metabarcoding approach to characterize both gut bacteria (16S: gut microbiome) and diet composition from foraged pollen (ITS2: floral diversity of bee pollen baskets), we aim to test the following predictions: (i) gut bacterial composition and diversity in *B. terrestris* will vary with environment (i.e. climate, land use), (ii) pollen packet diversity will correlate with gut bacterial diversity of *B. terrestris*, and show interactions with the environment. This is the first study in Australia to systematically investigate interactions among the gut microbial-dietary interface of an invasive pollinator and its novel environment, conducted at a landscape level.

## MATERIALS AND METHODS

### Study design and bumblebee collection

*B. terrestris* female workers were sampled from 19 sites across Tasmania, the Australian island state (Fig 1, Table 1), with site selection informed by prior records of the species occurrence (Hingston et al., 2002; Hingston, 2006). Bumblebees were typically collected from open areas of residential urban and rural regions, where flowering plants were present (such as road verges, flowery grass patches, coastal meadows, forest and national park margins, gardens, and parks), as described by Kardum Hjort et al. (2023a, b). During the active summer flight period in February 2020, free-flying *B. terrestris* workers were opportunistically captured from each site over a two-week period. The bumblebees were caught using handheld entomological sweep nets and jars, and the sampling was limited to 90 minutes. Captured worker bees were individually placed into 5ml plastic tubes, stored in a battery-powered car refrigerator (∼4°C) to induce chill coma. Subsequently, the bees were sexed (determined by presence of a stinger and or pollen packets) and placed in a freezer (– 18°C) for approximately 3 hours to induce euthanasia. Finally, the bees were preserved in 70% ethanol, following procedures outlined in Kardum Hjort et al. (2023a, b). For the gut microbiome study, we obtained data for 16/18 sites (n = 6 to 8 bees per site), while pollen samples were collected from 17/18 sites (pollen pooled from bees per site) (Fig 1, Table 1). Three sites (S8, S10 and S27) included in the pollenbiome study were not a part of the gut microbiome study because the bees from these sites had been previously damaged and their bodies not suitable for gut dissection. Conversely, two sites (S4 and S22) from the gut microbiome study were excluded from the pollenbiome study because the bees from these sites did not carry any pollen.

**Fig 1.**
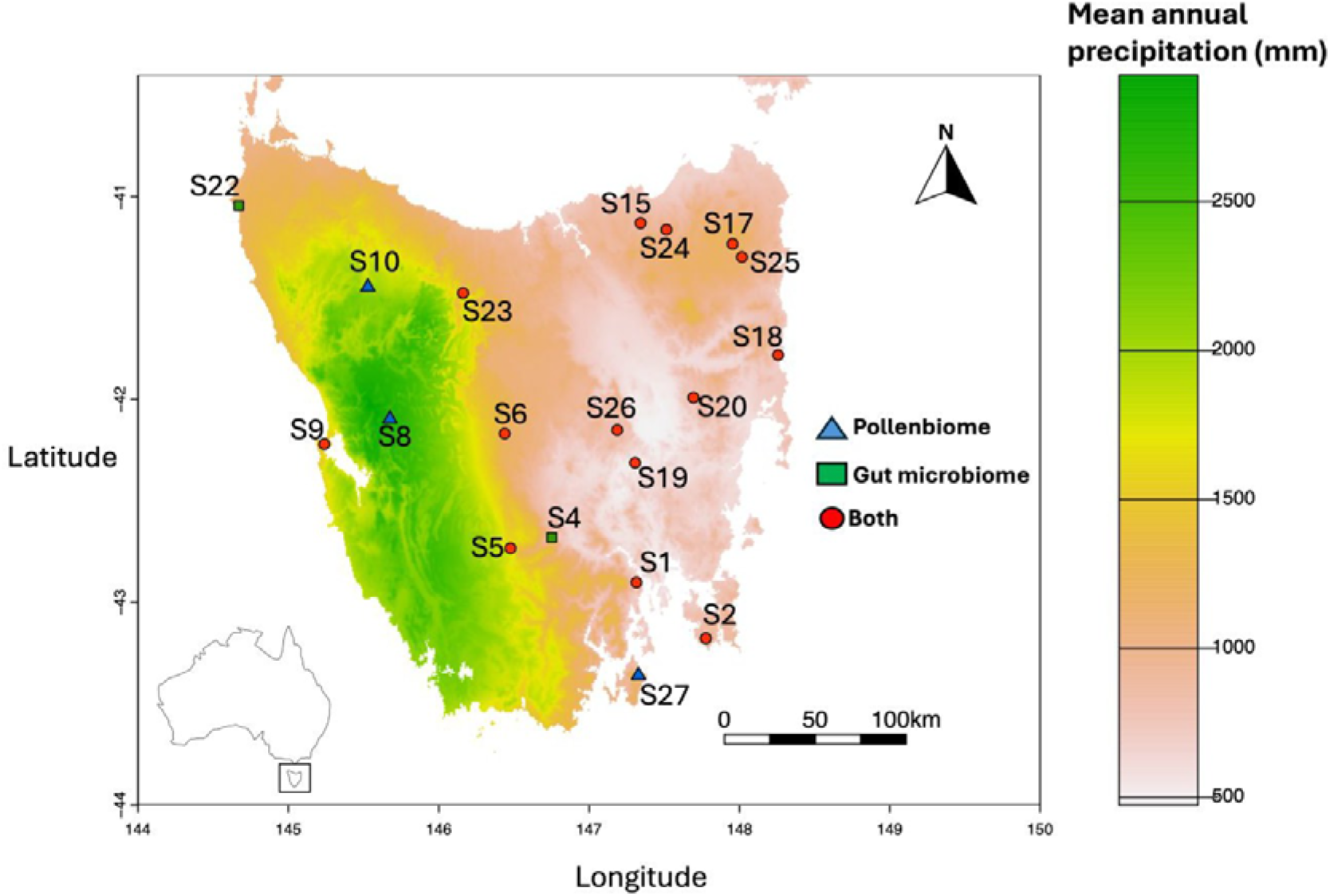
Sampling sites of *B. terrestris* (n=19) are shown (see Table 1 for site names) over mean annual precipitation (mm) across Tasmania, within Australia (inset). Key: ‘Pollenbiome’ = sites exclusively used in the pollenbiome study, ‘Gut microbiome’ = sites exclusively involved in the gut microbiome study, ‘Both’ = sites employed in both gut microbiome and pollenbiome studies.

**Table 1.**
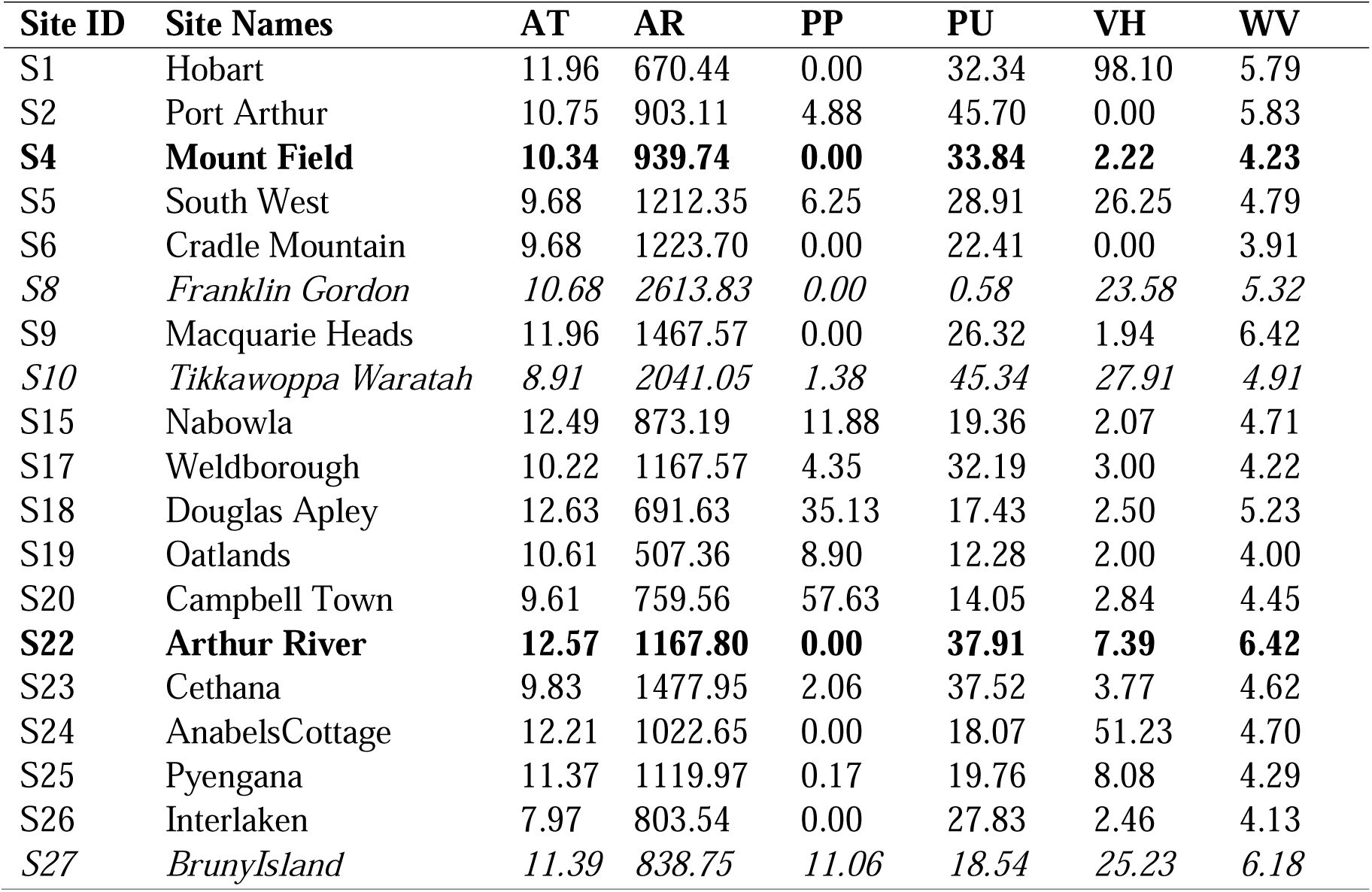
Environmental metadata extracted per sampling site. Sampling was conducted at a total of 19 sites across Tasmania, Australia. Cells emphasized in bold represent sites exclusively involved in the gut microbiome study (n=16), while the cells written in italics show sites exclusively used in the pollen biome study (n=17). All other sites were employed in both the gut microbiome and pollen biome studies. Abbreviations: AT = Mean annual temperature (□), AR = Mean annual precipitation (mm), PP = Percentage of pasture (%), VH = Height of vegetation (mm), PU = Percentage of urbanization (%), WV = Average velocity of summer wind (m/s).

### Selection of environmental variables

Our 19 sampling sites in Tasmania covered diverse climates, spanning the drier eastern to wetter western longitudes. The sites were sampled for environmental variables as described in Kardum Hjort (2023a, b) and summarised in Table 1. Environmental variables including mean annual temperature (□), mean annual precipitation (mm), precipitation seasonality (mm) and average summer wind velocity (m/s) were obtained from WorldClim v2.1 (Fick & Hijmans, 2017). Land cover details such as percentage of pasture was sourced from The National Dynamic Land Cover Dataset, while vegetation height (m) came from the ICESat Vegetation Height and Structure dataset (Scarth, 2011). The percentage of urban area was acquired from the Catchment Scale Land Use of Australia dataset (ABARES, 2021). These variables were extracted within a circular area around each site with a 1 km^2^ radius. Environmental variables were converted from cell fraction to the percentage of the total 1 km^2^ area (Kardum Hjort et al., 2023a; Kardum Hjort et al., 2023b).

### Environmental variables correlation

A Pearson’s correlation matrix was generated using R version 4.3.1 (R Core Team, 2024), to examine the relationships among all environmental variables across the sampled sites. Of all comparisons, only precipitation seasonality (mm) showed a high positive correlation (r > 0.7) with mean annual precipitation (mm) and was excluded from the analysis. The final set of predictor variables consisted of six environmental factors: mean annual temperature (□), mean annual precipitation (mm), percentage of pasture (%), height of vegetation (mm), percentage of urbanization (%), and average summer wind velocity (m/s) (Table S1).

### Pollen collection

The sample tubes containing *B. terrestris* workers and 70% ethanol were gently shaken to dislodge any pollen packets or grains attached to the bees and the bees were moved to a sterile petri dish. The collected pollen was transferred to a new Eppendorf tube, while the bee was returned to its original tube with ethanol. The pollen tube was centrifuged at 10000 rpm for three minutes. The ethanol supernatant was removed, ensuring removal of other debris while keeping the pollen pellet intact. The pollen pellet was washed in 200μl of DNase/RNase-free water (Invitrogen, Life Technologies) and the contents were transferred into a sterile 2ml ‘master tube’, which was centrifuged again at 10000 rpm for five minutes and the supernatant was removed. The procedure was repeated for each bee, with the pollen from each bee of the same site pooled within the master tube, which was then stored at −30□ until DNA extraction.

### Bumblebee gut dissection and gut bacterial DNA extraction

Gut dissections of the mid and the hindgut were performed for six to eight *B. terrestris* per site for 16 sites (Table S1). Prior to dissection, each *B. terrestris* individual was rinsed with 70% ethanol, placed on a small sterile petri dish, and immersed in sterile phosphate buffer solution (1x PBS, 137 mM NaCl; 2.7 mM KCl; 10 mM NaH_2_PO_4_; 1.8 mM KH_2_PO_4_). Dissections to separate the midgut and the hindgut from the body were carried out under a binocular stereo microscope (Motic SMZ 1711) with sterile forceps. The intact mid and hindgut were stored in 600μl 1xPBS. DNA was extracted from *B. terrestris* gut samples using a modification of the DNeasy Blood and Tissue Kit protocol (Qiagen). For pre-treatment of Gram-positive bacteria prior to the extraction, enzymatic lysis buffer (20 mM Tris-Cl, pH=8.0; 2mM Na_2_EDTA; 1.2% Triton X-100) was supplemented (20 mg ml^-1^) with lysozyme (Thermo Fisher Scientific), and 180 μl of this lysozyme-supplemented enzymatic lysis buffer was added to each gut sample. Glass beads (0.1 mm, Benchmark Scientific) were added to the samples and incubated on the heating block at 37□ for 45 minutes. The samples were homogenised using a Tissue Lyser II (Qiagen) for five minutes at 30Hz and then samples were incubated again at 37□ for 45 minutes. After lysis activity, 200μl of Buffer AL and 25μl of Proteinase-K (Ambion) was added to the samples and incubated at 56□ for 30 minutes. The rest of the steps for the DNA extraction were carried out based on the manufacturer’s protocol. DNA concentration was measured using a Qubit 4 Fluorometer using dsDNA High-Sensitivity (HS) Assay Kit (Invitrogen).

### Gut bacterial 16S rRNA metabarcoding and taxonomic identification

DNA metabarcoding of the *B. terrestris* gut microbiome was performed on the V4 region of the 16S rRNA gene for a total of 92 bumblebees, which was amplified using custom barcoded 515 forward primer (5’-TCGTCGGCAGCGTCAGATGTGTATAAGAGACAGCCTACGGGNGGCWGCAG-3’) and 806 reverse primer (5’-GTCTCGTGGGCTCGGAGATGTGTATAAGAGACAGGACTACHVGGGTATCTAATC C-3’) primers. The library preparation and sequencing were carried out at the Ramaciotti Centre for Genomics (University of New South Wales, Sydney, Australia) (Text S1), with a 2×250bp paired-end run on an Illumina MiSeq platform. Bioinformatics was performed on raw sequence data using Quantitative Insights into Microbial Ecology (QIIME-2, v2022.8, Bolyen et al., 2019). Demultiplexed paired-end reads were quality filtered using q2-demux plugin, followed by denoising with Deblur (Amir et al., 2017). All amplicon sequence variants (ASVs) were aligned with mafft, via q2-alignment (Katoh et al., 2002), and used to construct a phylogeny with fasttree2, via q2-phylogeny (Price et al., 2010). The generated ASVs were assigned to taxonomy using a pre-trained Naïve Bayes classifier against the Silva-138 reference database (Quast et al., 2012; Yilmaz et al., 2014; Bolyen et al., 2019) of the 515F/R806 region of the 16S rRNA gene and the q2-feature-classifier plugin (Bokulich et al., 2018; Robeson et al., 2021).

### Data filtering and statistical analysis of 16S rRNA data

Sequencing of the bacterial 16S rRNA amplicons yielded 9,679,597 reads (n=92 bumblebees from 16 sites). Following demultiplexing and quality filtering, a total of 6,396,272 reads were retained. Sequences shorter than 160 bp were excluded, and the sequence data were rarefied at a sequencing depth of 40,000 reads per sample, generating a total of 708 ASVs. ASVs exhibiting a relative abundance greater than 1% were determined using QIIME-2 to identify common gut bacterial species within bumblebees at each site (Estaki et al., 2020). To explore similarities in community composition across sites, a Bray-Curtis similarity matrix was generated using ASV abundance and visualised using a non-metric multidimensional scaling (NMDS) ordination plot which was constructed using the *vegan* R package (version 2.6-4) (Oksanen et al., 2024). The six environmental (predictor) variables were overlaid on the NMDS plot to examine their respective influence on the gut bacterial community composition of *B. terrestris*. Pairwise Permutational Multivariate Analysis of Variance (PERMANOVA), implemented using the *vegan* and *pairwise Adonis* R package (version 0.4.1) (Martinez Arbizu, 2020), was employed to detect significant variations in gut bacterial community composition among groups of sites. Chao1 richness and Shannon’s diversity indices of gut bacteria were computed using the *phyloseq* R package (version 1.44.0) (McMurdie & Holmes, 2013). Analysis of Variance (ANOVA) test and t-test were conducted between sites using *phyloseq* to assess for significant differences in Shannon’s diversity and Chao1 richness between sites, respectively.

### Pollen DNA extraction and ITS2 metabarcoding

The pooled pollen samples from each site were extracted for DNA using a modified protocol with the NucleoSpin Food Kit (Macherey Nagel). The CF (lysis) buffer was heated in a heating block for 10 minutes at 65□, and 1 ml of the heated buffer was added to each pollen sample. Glass and zirconium oxide beads (2 mm, Lysing Matrix H; MP Biomedicals) were added to each sample tube, which was then homogenised using a Tissue Lyser II (Qiagen) for three minutes at 20 Hz. Proteinase-K (30μl) was then added, and samples were incubated at 65□ in for 1.5 hours. The subsequent steps of the pollen DNA extraction adhered to the manufacturer’s protocol. Quantification of the extracted DNA was conducted using a Qubit 4 Fluorometer with the dsDNA HS Assay Kit (Invitrogen).

The Internal Transcribed Spacer region 2 (ITS2) is a variable DNA region in plants, spanning 100 to 700 bp, and is known for its remarkable discriminatory capabilities in pollen metabarcoding studies, particularly at the genus level (Yao et al., 2010; Milla et al., 2022). PCR was conducted on the extracted pollen DNA samples to amplify the ITS2 region, with 12.5μl of AmpliTaq Gold 360 MasterMix (Life Technologies), 0.5 μl of forward primer S2F (5’-ATGCGATACTTGGTGTGAAT-3’) (0.2 μM), 0.5 μl of reverse primer S3R (5’-GACGCTTCTCCAGACTACAAT-3’) (0.2 μM), 8.5 μl of DNase/RNase-free water (Invitrogen, Life Technologies), and 3μl of pollen DNA sample (Chen et al., 2010). The PCR protocol consisted of an initial denaturation at 94□ for 5 minutes, followed by 30 cycles of 94□ for 30 seconds, 56□ annealing for 30 seconds, 72□ extension for 45 seconds and a final extension at 72□ for 10 minutes. At the Ramaciotti Centre for Genomics (UNSW, Sydney, Australia), PCR products were subjected to purification, followed by a secondary PCR clean-up, library preparation and 2×250 bp paired-end sequencing was performed on an Illumina MiSeq platform (Text S1).

### Data filtering of pollen ITS2 data and plant taxa identification

A combined total of 1,100,167 forward and 1,100,167 reverse reads were obtained from pollen samples across our 17 sites. Primer sequences were identified and eliminated using *cutadapt* version 0.4 (Martin, 2011). The reads were subsequently processed using the *dada2* R package version 1.8, following the ITS pipeline workflow (Callahan et al., 2017).

Following an inspection of quality profiles, the reads were subjected to filtering and trimming according to standard parameters (maxN=0, maxEE=(2,2), truncQ=2, rm.phix=TRUE). A minimum sequence length of 50 (minLen=50) was enforced to exclude exceedingly short sequences. Post-merging of the forward and reverse reads was performed, and removal of chimeras, with 416,268 reads retained, and generated a table that consisted of 865 ASVs. ASVs with fewer than 10 observations were removed from the table, resulting in a final count of 445 ASVs. Basic Local Alignment Search Tool (BLAST), utilizing the *blastn* suite, was employed to assign genus to each DNA sequence from the ASV table. Euclidean distance matrix was used as a measure to explore the similarity among sites, relying on the sum of ASVs of plant genera per site.

### Interactions between gut bacteria, pollenbiome and environmental variables

Interactions of Shannon’s diversity indices of the gut microbiome and pollenbiome (response variables) with environmental factors (predictor variables) were analysed using the *lme* function in the *nlme* R package (version 3.1-166) (Pinheiro et al., 2024). Various linear mixed effect models were utilized with the *lme* function to investigate the: (1) impact of pollen diversity on gut bacterial diversity, (2) the influence of environmental variables on gut bacterial diversity, and (3) the combined interaction effects of environmental variables alongside pollen diversity on gut bacterial diversity. The models were then compared based on AIC using the *dredge* function within the *MuMIn* R package (version 1.47.5) (Bartoń, 2023). To achieve this, the percentage of pasture values first underwent logit transformation to linearize the relationship before conducting the *dredge* analysis. The sum of weights for the *dredge* models were calculated using the *sw* function in the *MuMIn* R package. Similarly, the interactions between Chao1 richness of gut bacteria, Chao1 richness of pollenbiome and environmental variables was also analysed using linear mixed effect models. However, comparisons based on AIC and sum of weights were not conducted between Chao1 richness indices of gut bacteria, pollenbiome and environment due to its lack of significance in the *lme* analysis (Table S2).

## RESULTS

### Gut bacterial taxa across sites

Proteobacteria, Firmicutes, and Actinobacteriota constituted the primary gut bacterial phyla in the bumblebee workers across all 16 sites (Fig 2A). Proteobacteria were the most abundant phyla in most sites, except for S9 (43.9%), where Firmicutes were prevalent (48.7%) (Fig 2A). A total of 16 major gut bacterial families were identified across the sites (Fig 2B). Among these, 12 families belonged to Proteobacteria, while three and one family were within the Firmicutes and Actinobacteriota phyla, respectively. The family *Pseudomonadaceae* was widespread across many sites. *Neisseriaceae* and *Morganellaceae* exhibited high relative abundances in S1 (28.4%) and S2 (33.72%) respectively, while *Lactobacillaceae* dominated in S9 (48.6%) and S22 (36.3%). Similarly, *Bifidobacteriaceae* was the most abundant bacterial family in S17 (27.4%) and S24 (32.7%) (Fig 2B). Analysis of site clusters, based on the relative abundance of major gut bacterial families, revealed distinct groupings among the sites (Fig 3).

**Fig 2.**
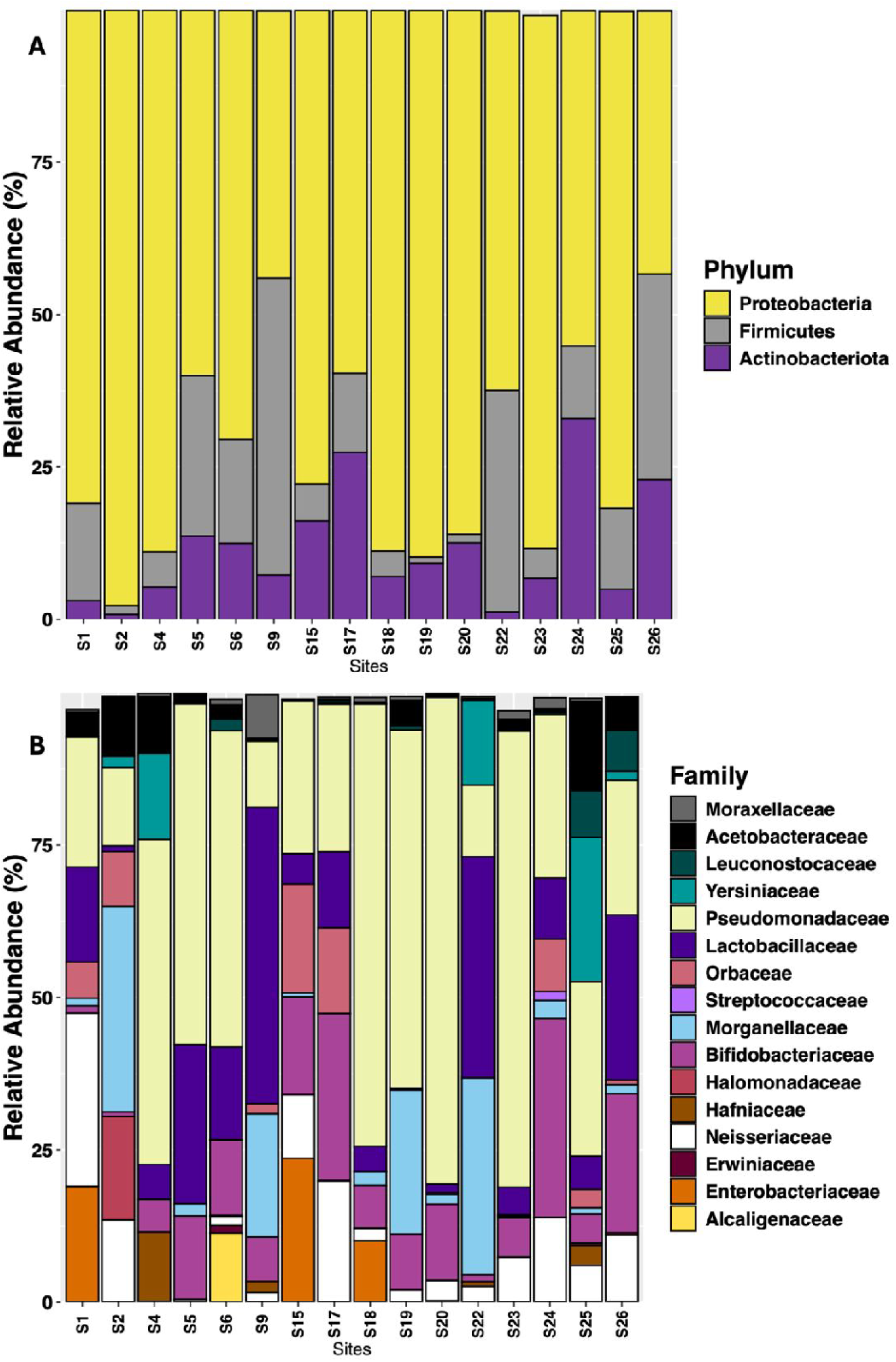
Percentage of relative abundance of *B. terrestris* gut bacteria sampled across 16 Tasmanian sites. (A) Relative abundance of core bacterial phyla per site. (B) Relative abundance of major bacterial families per site.

**Fig 3.**
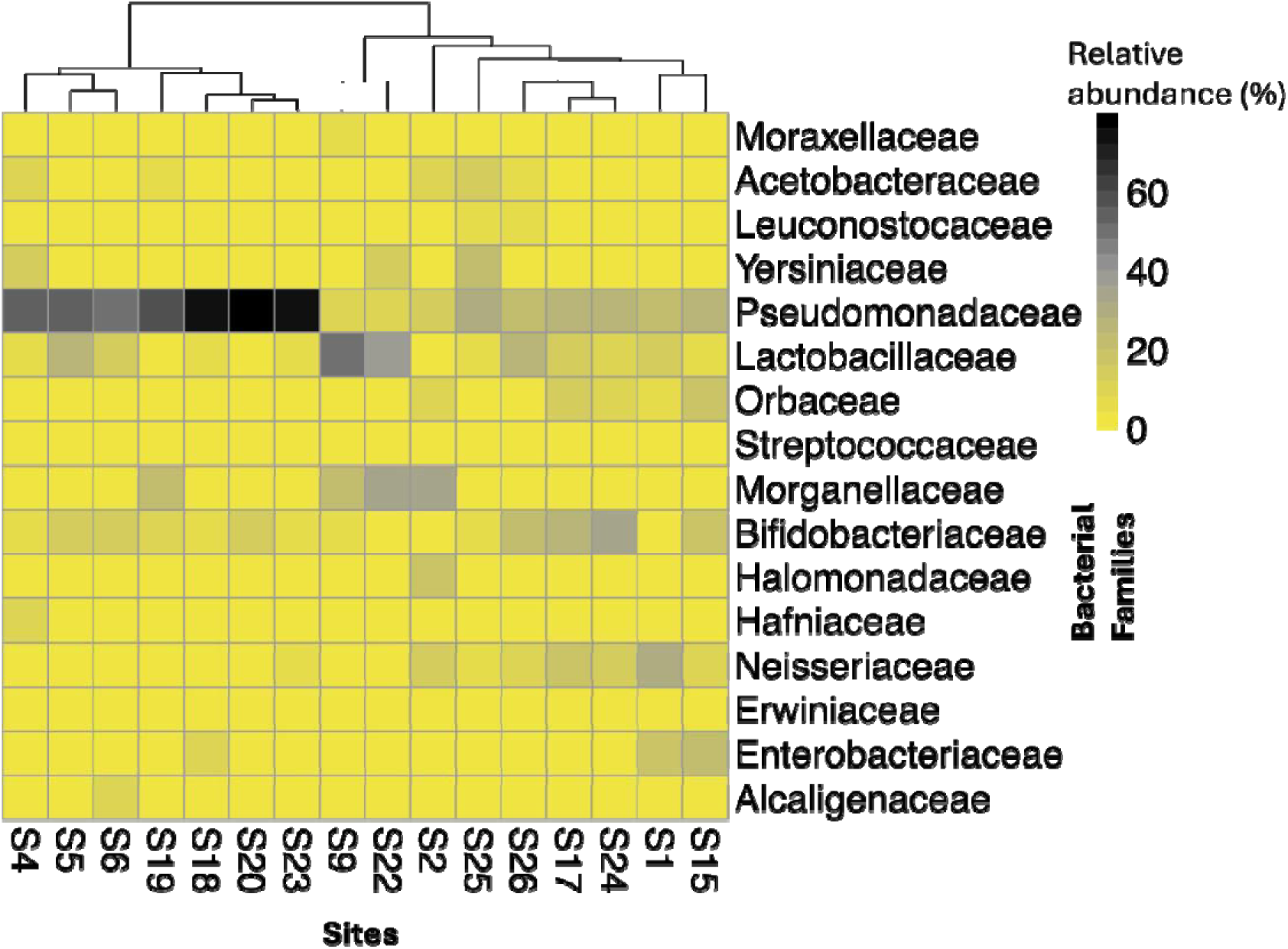
Heatmap showing the percentage relative abundance of major gut bacterial families within *B. terrestris* across each site. The colour scales show the relative abundance of different bacterial families within each site. The dendrogram on top of the heatmap depicts the distance or similarities among sites, which is constructed based on outcomes of the hierarchial clustering calculation using the Euclidean distance matrix, revealing the specific nodes to which each site is assigned.

The first cluster in the dendrogram was likely associated with the highest relative abundance of *Pseudomonadaceae*, encompassing sites S4 (53.3%), S5 (55.9%), S6(51.9%), S18 (72.6%), S19 (58.7%), S20 (79.8%) and S23 (74.8%). Apart from S5, S6 and S23, all the sites in the first cluster were located within the dry eastern region of Tasmania. In contrast, the second cluster grouped sites S1, S2, S9, S15, S17, S22, S24, S25 and S26 together, where *Pseudomonadaceae* contributed less than 50% to the overall gut bacterial community (Fig 3). The sites in the second cluster were dispersed throughout Tasmania without a distinct geographical pattern.

### Gut bacterial community composition and environmental variables

NMDS ordination with environmental variables showed that the percentage of pasture and mean annual precipitation were associated with gut bacterial community composition (Fig 4). The highest percentage of pasture was observed for S18 and S20 (S18=35.1%, S20=57.6%, Table 1), which appeared to drive the differences in the gut bacterial community structure, with both sites showing a relatively high abundance of *Pseudomonadaceae* (S18=72.5%, S20=79.9%, Fig 2B). Notably, S23 exhibited high relative abundance (74.8%) for *Pseudomonadaceae* (Fig 2B), yet no relationship was observed with percentage of pasture (2.1%). Sites S17 and S25 fell within an intermediate rainfall range (1168 mm and 1120 mm respectively) compared to other sites examined and appeared to drive the relationship between bacterial community composition and precipitation (Table 1).

**Fig 4.**
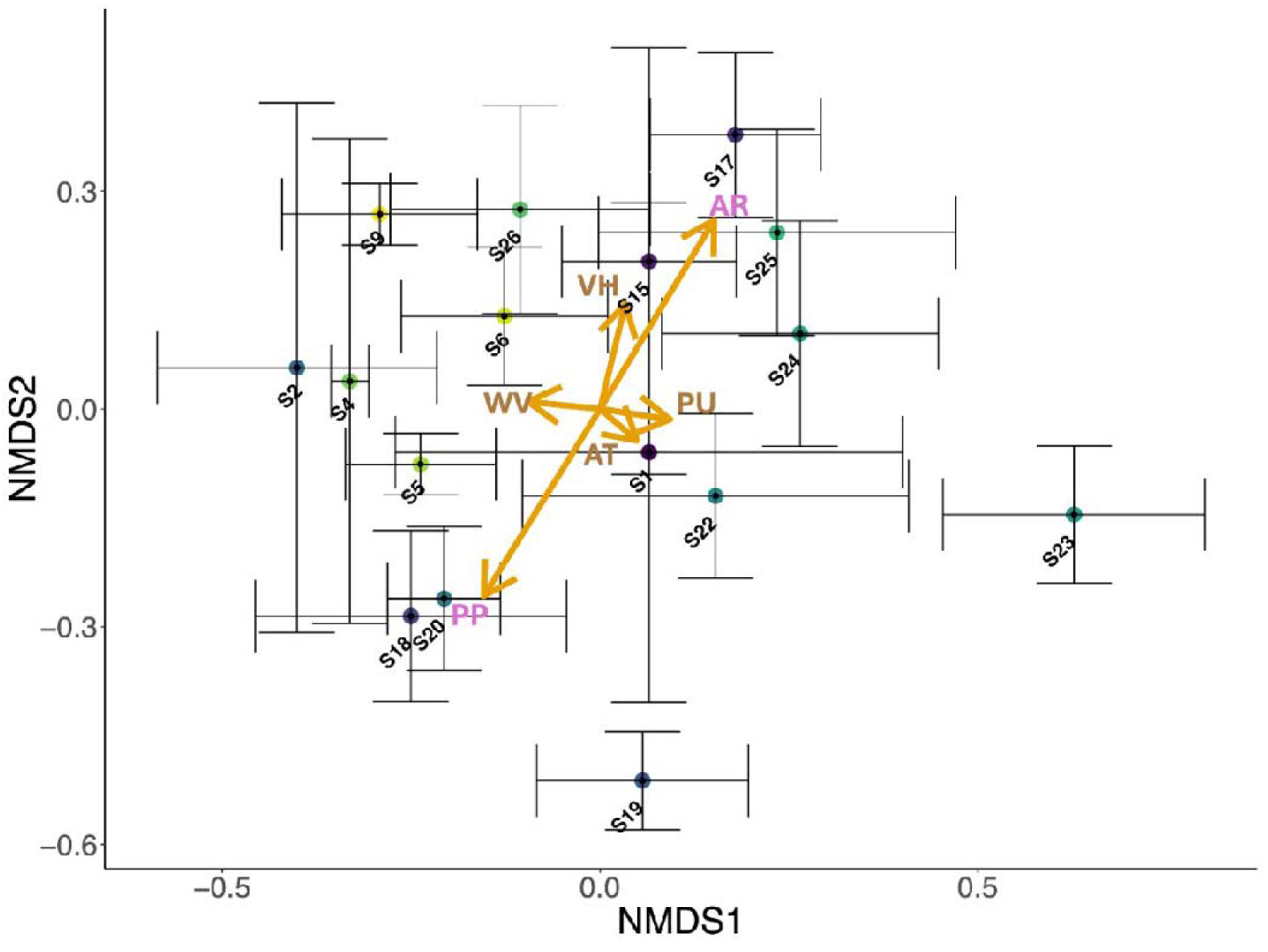
Ordination of gut bacterial communities for *B. terrestris* across 16 Tasmanian sites. The NMDS plot was constructed based on Bray-Curtis dissimilarity matrix of the amplicon sequence variant (ASV) abundance. Circles depict the centroid of ASV communities at each site, determined through the mean of the ASV abundance per site. The standard error bars indicate 95% confidence intervals for ordination axes. Greater proximity of the centroids signifies a higher similarity in the community composition of sites. The length of the arrows reflect the magnitude of impact of each environmental variable on the bacterial community composition. Abbreviations: AT = mean annual temperature (□), AR = mean annual precipitation (mm), PP = percentage of pasture (%), VH = height of vegetation (mm), PU = percentage of urbanization (%), and WV = velocity of summer wind (m/s).

In pairwise PERMANOVA, S9 displayed significant differences (p < 0.05) in bacterial community composition for 13 out of 16 sites (Table S3), excluding sites S5 and S22. Likewise, S22 showed significant differences (p < 0.05) for 12 out of 16 sites, excluding sites S1, S2 and S9 (Table S3). However, S9 and S22 were not significantly different from each other (p=0.194) (Table S3) and both had the highest prevalence of *Lactobacillaceae* (S9 = 48.6%, S22 = 36.3%, Fig 2B). The bumblebee gut bacteria at sites S23 and S25 had the greatest alpha diversity, using both Chao1 richness and Shannon’s diversity indices (Fig S1). The Chao1 richness index did not show significant differences in alpha diversity across all sites (*t-test*: p > 0.05, Fig S1A, Table S4). In contrast, the Shannon’s diversity index of S9 showed statistical significance with S19 and S20 (ANOVA: p < 0.05, Fig S1B, Table S5).

### Plant taxonomic composition

A total of 51 plant genera were detected from *B. terrestris* pollen packets across the 17 sites (Fig S2). Among these genera, 20 comprised of up to 99% of the total floral abundance (Fig 5A). When visualising site similarity in community structure in a heatmap-dendrogram, three clusters of sites could be distinguished by the cumulative ASV counts of plant genera (Fig S2). The first cluster of the dendrogram consisted of sites S17 and S18, which had the highest number of ASVs associated to *Rubus* spp. (19852 and 21554 ASVs respectively). The second cluster comprised of S1, S15, S19, S23, S26 and S27, which had the greatest number of ASVs attributed to *Eucalyptus* spp. (ranging from 9705 to 21274 ASVs). Except for S23, all sites from the first and second clusters were situated in the dry eastern part of Tasmania. The third cluster primarily consisted of sites with a higher prevalence of exotic plant genera or genera that have a combination of native and introduced species (Fig S2). The sites in the third cluster were spread across various parts of Tasmania, without a distinct geographical association. Among the 51 plant genera identified, 35 are introduced into Tasmania, 10 are endemic to Tasmania while 6 genera are known to contain both native and introduced species within Tasmania (Fig 5B, Table S6).

**Fig 5.**
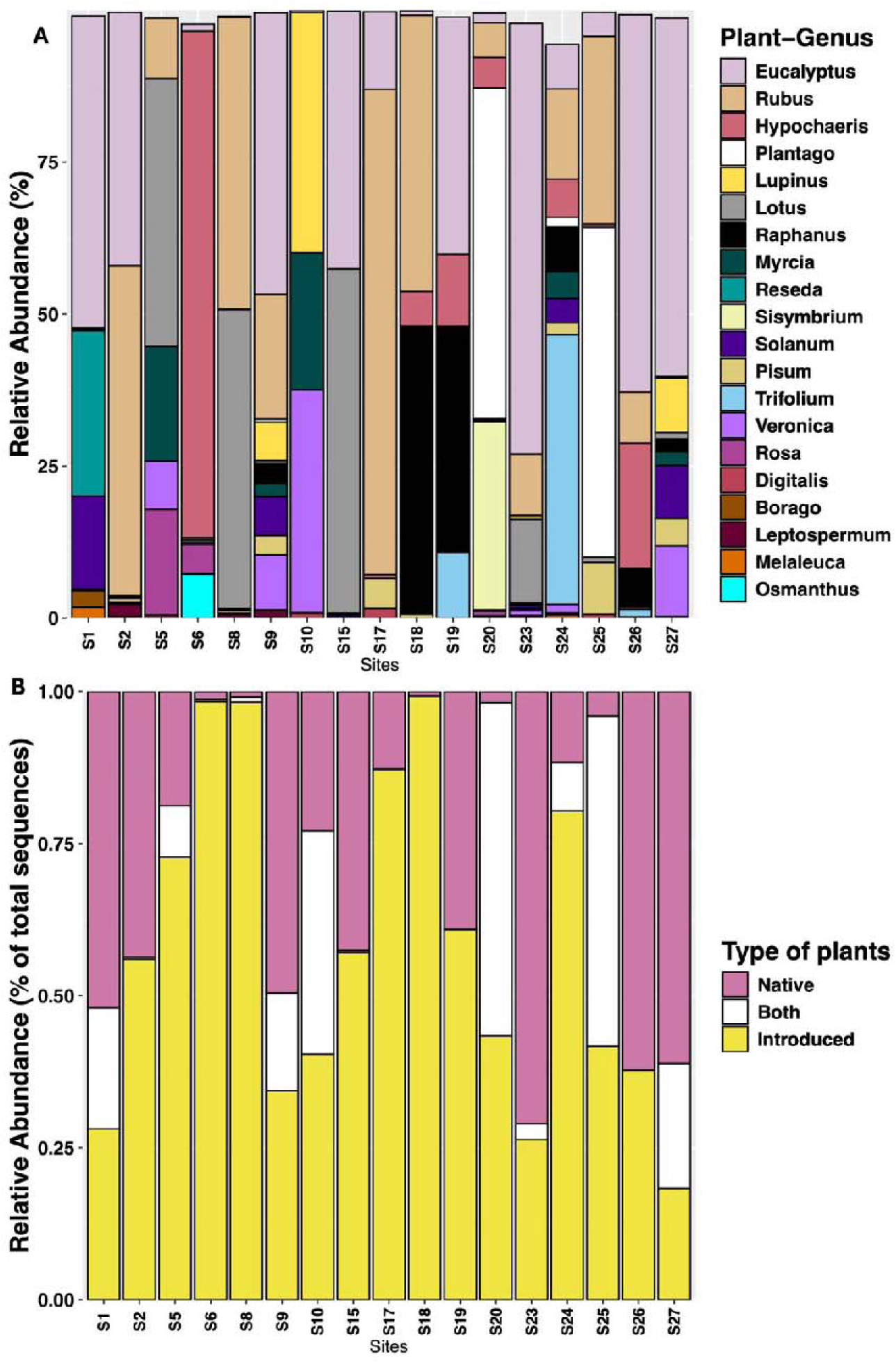
(A) Percentage of total sequences (relative abundance) of major plant genera identified from *B. terrestris* pollen packets across each sampling site. (B) Distribution of type of plants foraged by *B. terrestris* across each Tasmanian site. Explanation of type of plant category: ‘Native’ = plant genera that are native to Tasmania, ‘Introduced’ = plant genera that are exotic and have been naturalized in Tasmania, ‘Both’ = plant genera consisting of both native and introduced species in Tasmania (see Table S6 for plant genera within each category).

### Effects of pollen diversity and environment on gut bacterial diversity

Linear mixed-effect modelling revealed a significant positive interaction effect between mean annual precipitation and mean annual temperature on *B. terrestris* gut bacterial (Shannon’s) diversity (*lme*, DF=12, t=2.36, p=0.04, Fig 6A, Table 2). Further, there was a significant positive linear relationship between gut bacterial diversity and mean annual precipitation (*lm*, p=0.007, R^2^=0.4, Fig S3). There was also a positive interaction effect between average summer wind velocity and pollen (Shannon’s) diversity on gut bacterial diversity (*lme*, DF=10, t=2.76, p=0.02, Fig 6B, Table 2). However, the linear relationship between gut bacterial diversity versus pollen diversity and gut bacterial diversity versus average summer wind velocity were both statistically insignificant (*lm*, p > 0.05, Fig S4). Moreover, the relationship between pollen diversity and average summer wind velocity was statistically insignificant (*lm*, p=0.49, Fig S5D), and there were no other significant linear relationships between pollen diversity and any of the other environmental variables examined (Fig S5).

**Fig 6.**
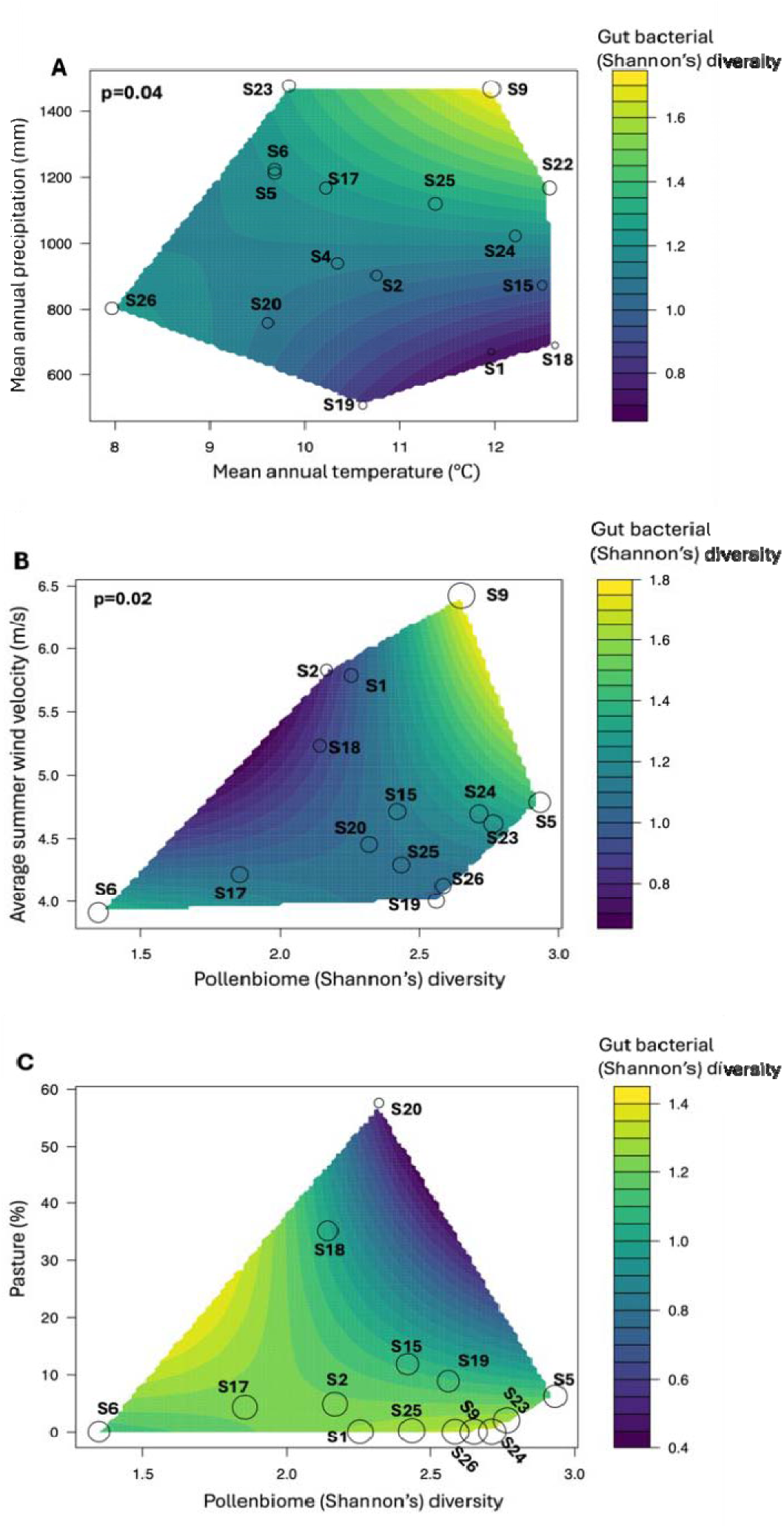
(A) Linear mixed effect model showing significant positive interaction effect of mean annual precipitation x mean annual temperature on gut bacterial diversity. (B) Linear mixed effect model showing significant positive interaction effect of pollen diversity x average summer wind velocity on gut bacterial diversity. (C) Relationship between percentage of pasture (most strongly weighted variable) and pollen diversity on gut bacterial diversity. In all three plots, the actual paired combinations of x-axis and y-axis used to fit the model are depicted as points with circle radii (sites) proportional to average gut bacterial (Shannon’s) diversity of *B. terrestris*, and the change in colour scale represents the change in gut bacterial diversity.

**Table 2.** Linear mixed-effect models and interaction effects between *B. terrestris* gut bacterial diversity, pollen packet diversity and environmental variables. Cells emphasized in bold show statistically significant interactions with p < 0.05. Abbreviations: Bacterial_diversity = Shannon’s diversity of gut bacterial samples, Pollen_diversity = Shannon’s diversity of pollenbiome samples, AT = Mean annual temperature (□), AR = Mean annual precipitation (mm), PP = Percentage of pasture (%), VH = Height of vegetation (mm), PU = Percentage of urbanization (%), and WV = Average velocity of summer wind (m/s).

| **Random effect ~1\|Sites** |  |  |  |
| --- | --- | --- | --- |
| **Fixed effect(s)** | **DF** | **t-value** | **p-value** |
| Bacterial_diversity~AT | 12 | -0.47 | 0.65 |
| **Bacterial_diversity~AR** | **12** | **2.87** | **0.01** |
| **Bacterial_diversity~PP** | **12** | **-2.44** | **0.03** |
| Bacterial_diversity~PU | 12 | 0.16 | 0.80 |
| Bacterial_diversity~VH | 12 | 0.58 | 0.57 |
| Bacterial_diversity~WV | 12 | 0.33 | 0.75 |
| **Bacterial_diversity~AT\*AR** | **10** | **2.36** | **0.04** |
| Bacterial_diversity~AT*PP | 10 | -0.43 | 0.67 |
| Bacterial_diversity~AT*PU | 10 | 0.28 | 0.78 |
| Bacterial_diversity~AT*VH | 10 | -0.75 | 0.47 |
| Bacterial_diversity~AT*WV | 10 | 1.47 | 0.17 |
| Bacterial_diversity~AR*PP | 10 | -0.58 | 0.58 |
| Bacterial_diversity~AR*PU | 10 | 0.39 | 0.70 |
| Bacterial_diversity~AR*VH | 10 | -1.90 | 0.08 |
| Bacterial_diversity~AR*WV | 10 | 0.99 | 0.35 |
| Bacterial_diversity~PP*PU | 10 | 0.26 | 0.80 |
| Bacterial_diversity~PP*VH | 10 | -1.01 | 0.33 |
| Bacterial_diversity~PP*WV | 10 | -0.46 | 0.65 |
| Bacterial_diversity~PU*VH | 10 | -1.27 | 0.23 |
| Bacterial_diversity~PU*WV | 10 | -1.16 | 0.27 |
| Bacterial_diversity~VH*WV | 10 | -2.09 | 0.06 |
| Bacterial_diversity~Pollen_diversity | 12 | 0.51 | 0.62 |
| Bacterial_diversity~Pollen_diversity*AT | 10 | 1.67 | 0.13 |
| Bacterial_diversity~Pollen_diversity*AR | 10 | -1.52 | 0.16 |
| Bacterial_diversity~Pollen_diversity*PP | 10 | -0.76 | 0.46 |
| Bacterial_diversity~Pollen_diversity*PU | 10 | 0.84 | 0.42 |
| Bacterial_diversity~Pollen_diversity*VH | 10 | 0.32 | 0.76 |
| **Bacterial_diversity~Pollen_diversity\*WV** | **10** | **2.76** | **0.02** |

When models were evaluated using Akaike information criterion (AIC), the top linear model (with a delta of 0, AIC of 131.6 and weight of 0.19) included the independent effects of percentage of pasture, pollen diversity and wind velocity, along with the interaction effects of pasture x pollen diversity and wind velocity x pollen diversity (Table S7). A comparison of the sum of weights further showed that percentage of pasture was the most strongly weighted variable (1.00) among all models that predicted gut bacterial diversity (Table 3). This was followed by pollen diversity (0.82) and the combination of both pasture and pollen diversity (0.77), wind velocity (0.64), the combination of pollen diversity and wind velocity (0.64) and mean annual precipitation (0.53). The remaining variables had much lower (< 0.13) AIC weightings (Table 3). The percentage of pasture exhibited a significant negative linear effect on bumblebee gut bacterial diversity (*lm*, p=0.03, R^2^=0.3, Fig S6). Upon examining the combined association of the three variables, it was observed that while the percentage of pasture increased, pollen diversity also increased, however, gut bacterial diversity decreased (Fig 6C). Notably, this relationship is largely driven by two specific sites, S18 and S20, which had the highest percentages of pasture (Fig 6C).

**Table 3.** Comparison of AIC sum of weights for the effect of pollen diversity (Shannon’s) and environmental variables on *B. terrestris* gut bacterial diversity(Shannon’s). The most strongly weighted variable is emphasized in bold. Abbreviations: logit_pasture = logit transformed values for percentage of pasture, pollen_diversity = Shannon’s diversity for pollenbiome samples, wind = average summer wind velocity (m/s), precipitation = mean annual precipitation (mm).

| Variable | sum of weights | N containing models |
| --- | --- | --- |
| **logit_pasture** | **1** | **9** |
| pollen_diversity | 0.82 | 7 |
| logit_pasture : pollen_diversity | 0.77 | 6 |
| wind | 0.64 | 4 |
| pollen_diversity : wind | 0.64 | 4 |
| precipitation | 0.53 | 7 |
| logit_pasture : wind | 0.12 | 1 |
| precipitation : logit_pasture | 0.12 | 2 |
| precipitation : pollen_diversity | 0.08 | 1 |

## DISCUSSION

The health of bees is closely linked to their nutrition and the composition of their gut microbiomes, yet the influence of natural environmental conditions on bee health remains understudied, particularly in invasive contexts. Here, we find that increased pollen diversity in sites with an increased percentage of pasture is associated with reduced gut bacterial diversity in the recently introduced *B. terrestris* on the Australian island state of Tasmania (Fig 6C). In contrast, rainfall positively affected the diversity of gut bacteria in *B. terrestris*, whether considered alone or in combination with increasing temperatures (Fig 6A, Fig S3). Additionally, sites with higher average summer wind velocities and higher pollen diversity showed a positive association with the gut bacterial diversity of *B. terrestris* (Fig 6B). As expected, introduced plant genera dominated the pollen taxa (Fig 5B, Table S6). Furthermore, both the percentage of pasture and mean annual precipitation were identified as significant factors influencing the gut bacterial community composition (Fig 4). Our findings shed light on how the interplay between an invasive pollinator and its novel environment impacts pollinator gut bacterial diversity and composition. This contributes to understanding factors influencing the distribution and persistence of invasive pollinators in novel landscapes.

### Environmental effects on B. terrestris gut bacteria

We observed varying degrees of significant differences among the sites in our gut microbiome study (n=16) based on the composition of the *B. terrestris* gut bacterial community, as indicated by pairwise PERMANOVA (Table S3). Notably, sites S9 and S22 exhibited the most significant differences with other sites (Table S3). However, there was no significant difference between S9 and S22 themselves. These observations could be attributed to both S9 and S22 being more isolated on the west coast of Tasmania. Further, it was previously found that these two sites showed genetic differentiation from all other sites sampled across Tasmania (Kardum Hjort et al., 2023a). The findings of this study indicated that the percentage of pasture can predict the gut bacterial community composition (Fig 4) and negatively impact the diversity of gut bacteria in *B. terrestris* across Tasmania (Fig S6). Sites with the highest percentage of pasture were positioned in the northern midlands (S20) and in a national park locality within the east coast (S18) of Tasmania. These areas are characterized by intensive pasture-based agricultural practices linked to high-intensity grass cropping that leaves few floral species (Lane et al., 2015). Notably, a previous study exploring the microbiome of bee bread (stored pollen) from honeybees and its relationship with the environment suggested that improved grasslands, with reduced floral diversity, lead to a decrease in gut bacterial community structure and diversity (Donkersley et al., 2018). Previous research in Tasmania also revealed that the proboscis length of *B. terrestris* was reduced in areas with a high percentage of pasture (Kardum Hjort et al., 2023b). This latter observation supports possible selection pressures on proboscis length in pastural areas due to differing floral characteristics of pastural plants (e.g., shorter floral tubes).

Mean annual precipitation was associated with the community composition of bumblebee gut bacteria (Fig 4). The two sites that appeared to be driving this association, sites S17 and S25, were in the north-eastern region of Tasmania, with an intermediate range of mean annual precipitation (Hill et al., 2009). We found that an increase in mean annual precipitation significantly increased gut bacterial diversity (Fig S3), while sites with higher mean annual temperature and higher mean annual precipitation, thus higher humidity, also had a positive effect on gut bacterial diversity (Fig 6A). Precipitation and humidity have a strong impact on insect distributions (Koot et al., 2022) and physiological processes, such as regulation of water content within their bodies (Siepielski et al., 2017; Singh et al., 2009). Bees are particularly sensitive to reduced precipitation and drier conditions (Jackson et al., 2018). Research on the seasonal dynamics of the honeybee gut microbiota under subtropical climates suggested that bees undergo significant changes in their daily routines on cold and rainy days, as they cannot fly to collect nectar or pollen and tend to accumulate inside their colonies during such weather conditions (Castelli et al., 2022). Bees depend on visual information to navigate their surroundings, and precipitation can alter their spatial awareness, thus distorting their perception of the environment during foraging (Lawson & Rands, 2019). Such changes in high precipitation environments may therefore influence foraging activity and diet, hence altering individual bee gut microbiomes.

Notably, foraged pollen harbors a rich bacterial diversity that may shift with temperature or humidity (McFrederick and Rehan 2022), which can reduce pollen viability (Iovane et al. 2022) and change gut microbes. McFrederick and Rehan (2022) conducted a study of carpenter bees (*Ceratina australensis*) in Australia and found that the pollen microbiome differed in subtropical, temperate and grassland zones, suggesting that variation in pollen usage in bees across landscapes results in variation in ingested microbes. As the pollen microbiome can be transmitted to bees during foraging, this can translate to shared pollen and gut microbiomes in bees (Keller et al. 2021). These shared microbes may provide benefits to bees via detoxification or enhancing nutritional quality (Keller et al. 2021). Notably, Kardum Hjort et al., (2023a) showed that *B. terrestris* in Tasmania shows evidence for local adaptation in relation to precipitation and identified candidate genes under selection with functions related to cuticle-regulated water loss prevention and olfactory systems. Candidate genes under selection associated with precipitation have also been found in *Bombus vancouverensis*, and were also related to cuticle structure and function, which plays a role in desiccation tolerance (Jackson et al., 2020). This suggests that precipitation may affect both physiological and behavioural processes that impact evolutionary processes that include biotic interactions with bacterial communities.

We found that the positive interaction effect between average summer wind velocity and pollen diversity was significantly associated with an increase in bumblebee gut bacterial diversity (Fig 6B). The mixed interaction model recognized site S9 as having the highest gut bacterial diversity and the greatest average summer wind velocity (Fig 6B, Table 1). However, the observed insignificant linear relationships of gut bacterial diversity with pollen diversity (Fig S4A), gut bacterial diversity with average summer wind velocity (Figure S4B) and pollen diversity with average summer wind velocity (Figure S5D) suggest that other factors may be at play. It is possible that the remote location of S9, along with its proximity to the Southern Ocean and Macquarie Harbour, could be influencing these interaction effect. Wind is known to affect various aspects of bumblebee behaviour such as flight performance (Lawson & Rands, 2019; Chang et al., 2016), flight stability (Ravi et al., 2016), foraging energy expenditure (Crall et al., 2017), guidance to flower rewards (Kantsa et al., 2019) and decisions that involved choosing between pollen and nectar foraging during windy conditions (Mountcastle et al., 2015). A previous study found selection signatures within *B. terrestris* in Tasmania in relation to average summer wind velocity (Kardum Hjort et al., 2023a). Hence, environmental factors such as pasture, precipitation and wind appear to play important roles in determining both bee gut bacterial compositions and selection processes at the population level.

### Diversity and functions of prevalent gut bacterial taxa

Our study found that the gut bacterial samples of *B. terrestris* were predominantly composed of Proteobacteria, Firmicutes and Actinobacteriota (Fig 2A), aligning with findings from prior studies (Krams et al., 2022; Tang et al., 2023). These three bacterial phyla are prevalent not only in the European honeybee (*A. mellifera*) but also in various other bumblebee species, including *Bombus auricomus, Bombus bimaculatus, Bombus fervidus, Bombus griseocollis, Bombus impatiens, Bombus pensylvanicus* and *Bombus perplexus* (Kakumanu et al., 2016; Amiri et al., 2023; Rudra Gouda et al., 2024). Furthermore, these phyla are part of the gut microbiomes of Australian native bees like *Tetragonula carbonaria* and *Austroplebea australis* (Liu et al., 2023). At the family level, *Pseudomonadaceae* was widespread across most of the sites under the study (Fig 2B). The presence of diverse members of *Pseudomonadaceae* family is proposed to contribute to high metabolic adaptability, secreting a wide range of bioactive enzymes and secondary metabolites (for e.g., antimicrobial peptides, surfactants, siderophores and cell-wall-degrading enzymes) (Tsadila et al., 2023). Sites S9 and S22, which were located on the west coast of Tasmania, had the highest prevalence of *Lactobacillaceae* (Fig 2B). *Lactobacillaceae* has been recognized for its role in influencing learning and memory behaviour in *A. mellifera* through the regulation of tryptophan metabolism (Zhang et al., 2022). Similarly, bacterial members from *Bifidobacteriaceae* and *Neisseriaceae* (Fig 2B) play essential roles as primary decomposers of polysaccharides and contributors to innate immunity, respectively in the gut of honeybees (Bleau et al., 2020; Chen et al., 2021).

### Pollen diversity, nutrition and the bee gut microbiome

There have been reports of global large-scale colony losses of bees in recent years (Neumann & Carreck, 2010; Kulhanek et al., 2017; Requier et al., 2018; Gray et al., 2019). These losses have been linked with infection by multiple pests and pathogens, pesticide intoxication and depreciating food resources (Naug, 2009; Potts et al., 2010; Goulson et al., 2015; Barron 2015; Steinhauer et al., 2018). Two important aspects that can enhance the ability of bees to cope with colony-loss related challenges involve proper nutrition (Dolezal & Toth; 2018; Stanimirović et al., 2019) and functional worker gut microbiota (Alberoni et al., 2016; Kwong & Moran, 2016; Bonilla-Russo & Engel, 2018). Foraging bees rely exclusively on pollen for essential proteins, lipids, vitamins and minerals (Haydak, 1970; Herbert et al., 1978; Wright et al., 2018) that are vital for their overall growth and development (Winston, 1991; Brodschneider & Crailsheim, 2010). Digestion of pollen is complex (Nicolson et al., 2018), with the gut bacteria of bees playing a critical role in breaking down complex cell wall polysaccharides such as hemicellulose and pectin, which is essential for accessing the nutrients within pollen (Lee et al., 2015, Lee et al., 2018; Zheng et al., 2019). Disruptions within the gut microbiota have been linked to diseases and malnutrition (Lozupone et al., 2012), including reduced efficiency in honeybee protein digestion (du Rand et al., 2020).

*B. terrestris* is known for its generalist feeding habit, as it reportedly forages on both native and introduced plants (Hanley & Goulson, 2003; Hingston, 2007). Our study found that the foraged pollen taxa collected from the invasive *B. terrestris* population in Tasmania was dominated mainly by introduced plants (Fig 5B, Table S6). It was noted that despite being native, *Eucalyptus* emerged as a highly favoured diet among bumblebees across most Tasmanian sites (Fig 5A, Fig S2). Pollen from *Eucalyptus* spp. is characterized by a protein content of around 20% or less during the flowering period (Invernizzi et al., 2011; Branchiccela et al., 2019). *Eucalyptus* pollen has the lowest lipid content (Roulston & Cane, 2000; Branchiccela et al., 2019). Both pollen and lipid are required for colony maintencance and brood rearing (Branchiccela et al., 2019). It also contains omega-3 fatty acids (Arien et al., 2015) and essential amino acids, such as isoleucine (Somerville, 2001). Essential amino acids are required for growth and development of honeybees (Roulston et al., 2000; Cook et al., 2003), while omega-3 deficiency impairs honeybee learning (Arien et al., 2015). *Rubus* was identified as the most important introduced plant genus across most of the sites (Fig 5A, Fig S2). Pollen from various *Rubus* spp. consists of protein ranging from 15% to 37% (Vanderplanck et al., 2014; Conti et al., 2016; Liolios et al., 2016; Kostryco & Chwil, 2021) and amino acid between 32.8% and 35% (Hanley et al., 2008; Di Pasquele et al., 2013). The primary fatty acids in *Rubus* spp. pollen were α-linolenic, linoleic and palmitic acids (Feás et al., 2012). Saa-Otero et al. (2000) showed the presence of 19 fatty acids within bee pollen baskets of *Rubus* pollen grains. The fatty acids included linolenic acid, isopalmitic acid, palmitic acid, oleic acid and linoleic acid. Pollen with high fatty acid content may help to inhibit growth of infectious microorganisms (for e.g. *Paenibacillus* larvae, *Melissococcus plutonius*) (Manning, 2001). Certain lipids may act as ligands for allergenic proteins (Dahl, 2018) and play key roles in plant chemotaxonomy (Kostryco & Chwil, 2021). Therefore, our study emphasizes the importance of a diet containing pollen from various plant species in *B. terrestris,* as floral resources may be selected based on nutritional needs.

### Introduced bees adapting to novel diets

Since introduced bees typically prefer foraging on introduced plants, they often act as the primary pollinators for several weeds (Lowenstein et al., 2019, O’Connell et al., 2021). Our study revealed that out of 51 plant genera identified from pollen baskets of *B. terrestris* in Tasmania (Fig S2), the majority (n = 35 genera) were introduced plant genera, 10 were native plant genera, and six genera belonged to plants with both native and introduced species (Fig 5B, Table S6). In Tasmania, several exotic plants such as *Lupinus, Trifolium, Raphanus* remained scarce for many decades until the introduction of *B. terrestris* (Hanley & Goulson, 2003). In our study, we identified several invasive weeds such as *Hypochaeris, Lupinus, Trifolium, Sisymbrium, Digitalis, Cirsium* (Fig S2). The distribution and abundance of some introduced bees is closely tied to the availability of alien food plants. For instance, populations of *Bombus ruderatus* and *Bombus subterraneus* in New Zealand experienced significant declines following changes in agricultural practices that reduced the distribution of *Trifolium pratense* and *Lotus corniculatus* (Macfarlane & Gurr, 1995; Hanley & Goulson, 2003). However, the generalist, *B. terrestris,* exhibits a much more generalist foraging habit compared to many other introduced bee species, allowing them to feed on a variety of non-native and native plants (Butz Huryan, 1997; Hingston et al., 2002; Goulson 2003).

Therefore, while we find that *B. terrestris* derives advantages from the presence of alien weed species in Tasmania, their spread and persistence is unlikely to be hindered by the absence of introduced plants (Hanley & Goulson, 2003), given their use of native species also.

## Conclusion

Our study marks the first landscape-level investigation of the combined interplay between the gut microbiome and foraged pollen in an invasive bee pollinator following a recent (∼30-year-old) invasion into a novel environment. Our findings set a foundation for integrating biotic and abiotic processes into the management of both invasive pollinators and those requiring conservation. We acknowledge that the data represent a snapshot in time and therefore could be improved with temporal sampling of both pollen and gut microbiomes throughout the foraging period. A more comprehensive reference database of sequences for Tasmanian plants at higher taxonomic resolution would further improve our findings. This research sets the stage for future comparative analyses of invasive bumblebee gut microbiome and pollenbiomes with honeybees and native Australian bees, which may improve understanding of how bee gut microbiomes are shared across diverse landscapes, and their relationship to immunity and fitness. Here, we established how environmental interactions influence the gut microbiome of a pollinator in an invaded landscape with novel nutritional resources, which may help to predict where invasive pollinators can spread and persist.

## Acknowledgements

We would like to thank Emily Petrolo, Sanne Nielson who helped to collect samples in Tasmania, and Andrew Allen who provided valuable statistical advice. This project was funded by an Australian Research Council Future Fellowship awarded to R. Dudaniec (FT230100478), a Macquarie University Research Acceleration Scheme Grant (to R. Dudaniec) and the School of Natural Sciences at Macquarie University graduate student funding (to S. Haque). Permits for collecting *B. terrestris* in Tasmania were obtained from the Department of Primary Industries, Parks, Water and Environment, Tasmania (Authority No. FA 19253). A permit for transportation of deceased *B. terrestris* to New South Wales was obtained from the Department of Primary Industries, NSW Government (Ref: OUT19/15645).

## Author contributions

R. Dudaniec conceived the ideas and designed methodology; C. Kardum Hjort collected the samples and S. Haque and R. Dudaniec conducted laboratory work and generated genetic data; S. Haque undertook data analysis in consultation with H. Gamage and R. Dudaniec; S. Haque and R. Dudaniec led the writing of the manuscript. All authors contributed to data interpretation. All authors edited and contributed critically to the drafts and gave final approval for publication.

## Data availability statement

All datasets and code will be available on the Dryad Digital Repository upon acceptance of the article.

## References

Abares, 2021. Catchment Scale Land Use of Australia – Commodities – Update December 2020. Australian Bureau of Agricultural and Resource Economics and Sciences, Canberra, February CC BY 4.0. 10.25814/jhjb-c072

Abrol DP. (2012). Consequences of introduced honeybees upon native bee communities. In: Pollination Biology: Biodiversity Conservation and Agricultural Production. Springer. 635–658. 10.1007/978-94-007-1942-2

Acosta AL, Giannini TC, Imperatriz-Fonseca VL et al., (2016). Worldwide alien invasion: a methodological approach to forecast the potential spread of a highly invasive pollinator. Public Library of Science, 11(2), e0148295. 10.1371/journal.pone.0148295

Aizen MA, Arbetman MP, Chacoff NP et al., (2020). Invasive bees and their impact on agriculture. Advances in Ecological Research. 63, 49–92. 10.1016/bs.aecr.2020.08.001

Alaux C, Dantec C, Parrinello H et al., (2011). Nutrigenomics in honey bee: digital gene expression analysis of pollen’s nutritive effects on healthy and *Varroa*-parasitized bees. BMC Genomics, 12, 496. 10.1186/1471-2164-12-496

Alberoni D, Gaggìa F, Baffoni L et al., (2016). Beneficial microorganisms for honey bees: problems and progresses. Applied Microbiology and Biotechnology, 100(22), 9469–9482. 10.1007/s00253-016-7870-4

Amir A, McDonald D, Navas-Molina JA et al., (2017). Deblur rapidly resolves single-nucleotide community sequence patterns. MSystems, 2(2), e00191–16. 10.1128/mSystems.00191-16

Amiri N, Keady MM and Lim HC (2023). Honey bees and bumble bees occupying the same landscape have distinct gut microbiomes and amplicon sequence variant-level responses to infections. PeerJ – the Journal of Life and Enviromental Sciences, 11, e15501. 10.7717/peerj.15501

Anderson KE, Sheehan TH, Eckholm BJ et al., (2011). An emerging paradigm of colony health: microbial balance of the honey bee and hive (*Apis mellifera*). Insectes Sociaux, 58(4), 431–444. 10.1007/s00040-011-0194-6

Arien Y, Dag A, Zarchin S et al., (2015). Omega-3 deficiency impairs honey bee learning. Proceedings of the Natural Academy of Sciences, 112(51), 15761–15766. 10.1073/pnas.1517375112

Arstingstall KA, DeBano S, Li X et al., (2021). Capabilities and limitations of using DNA metabarcoding to study plant-pollinator interactions. Molecular Ecology, 30, 5266–5297. 10.1111/mec.16112

Baniel A, Amato KR, Beehner JC et al., (2021). Seasonal shifts in the gut microbiome indicate plastic responses to diet in wild geladas. Microbiome, 9, 26. 10.1186/s40168-020-00977-9

Barascou L, Sene D, Barraud A et al., (2021). Pollen nutrition fosters honeybee tolerance to pesticides. Royal Society of Open Science, 8, 210818. 10.1098/rsos.210818

Barron AB. (2015). Death of the bee hive: understanding the failure of an insect society. Current Opinion in Insect Science, 10, 45–50. 10.1016/j.cois.2015.04.004

Bartoń, K. (2023). MuMIn: Multi-model inference (R package version 1.47.5). 10.32614/CRAN.package.MuMIn

Bell KL, Fowler J, Burgess KS et al., (2017). Applying pollen DNA metabarcoding to the study of plant-pollinator interactions. Applications in Plant Sciences, 5(6), apps 1600124. 10.3732/apps.1600124

Bell KL, Turo KJ, Lowe A et al., (2022). Plants, pollinators, and their interactions under global ecological change: The role of pollen DNA metabarcoding. Molecular Ecology, 32(23), 6345– 6362. 10.1111/mec.16689

Biesmeijer JC, Roberts SPM, Reemer M et al., (2006). Parallel declines in pollinators and insect-pollinated plants in Britain and the Netherlands. Science, 313(5785), 351–4, 10.1126/science.1127863

Billiet A, Meeus I, Van Nieuwerburgh F et al., (2016). Impact of sugar syrup and pollen diet on the bacterial diversity in the gut of indoor-reared bumblebees (*Bombus terrestris*). Apidologie, 47, 548–560. 10.1007/s13592-015-0399-1

Bleau N, Bouslama S, Giovenazzo P et al., (2020). Dynamics of the Honeybee *(Apis mellifera)* Gut Microbiota Throughout the Overwintering Period in Canada. Microorganisms, 8(8), 1146, 10.3390/microorganisms8081146

Bokulich NA, Kaehler BD, Rideout JR et al., (2018). Optimizing taxonomic classification of marker-gene amplicon sequences with QIIME 2’s q2-feature-classifier plugin. Microbiome, 6, 90. 10.1186/s40168-018-0470-z

Bolyen E, Rideout JR, Dillon MR et al., (2019). Reproducible, interactive, scalable and extensible microbiome data science using QIIME 2. Nature biotechnology, 37, 852–857. 10.1038/s41587-019-0209-9

Bonilla-Rosso G and Engel P. (2018). Functional roles and metabolic niches in the honey bee gut microbiota. Current Opinion in Microbiology, 43, 69–76. 10.1016/j.mib.2017.12.009

Branchiccela B, Castelli L, Corona M et al., (2019). Impact of nutritional stress on the honeybee colony health. Scientific Reports, 9(1), 10156. 10.1038/s41598-019-46453-9

Brodschneider R and Crailsheim K. (2010). Nutrition and health in honey bees. Apidologie 41, 278–294. 10.1051/apido/2010012

Brodschneider R, Kalcher-Sommersguter E, Kuchling S et al., (2021). CSI Pollen: Diversity of Honey Bee Collected Pollen Studied by Citizen Scientists. Insects, 12(11), 987. 10.3390/insects12110987

Butz Huryn VM. (1997). Ecological impacts of introduced honeybees. The Quarterly Review of Biology, 72, 3, 275–297. 10.1086/419860

Callahan B, McMurdie P, Rosen M et al., (2017). DADA2: High-resolution sample inference from Illumina amplicon data. Nature Methods, 13, 581–583. 10.1038/nmeth.3869

Castelli L, Branchiccela B, Romero H et al., (2022). Seasonal Dynamics of the Honey Bee Gut Microbiota in Colonies Under Subtropical Climate: Seasonal Dynamics of Honey Bee Gut Microbiota. Microbial Ecology, 83(2), 492–500. 10.1007/s00248-021-01756-1

Chang JJ, Crall JD and Combes SA. (2016). Wind alters landing dynamics in bumblebees. Journal of Experimental Biology, 219(18), 2819–2822. 10.1242/jeb.137976

Chen J, Wang J and Zheng H. (2021). Characterization of *Bifidobacterium apousia* sp. nov., *Bifidobacterium choladohabitans* sp. nov., and *Bifidobacterium polysaccharolyticum* sp. nov., three novel species of the genus *Bifidobacterium* from honey bee gut. Systematic and Applied Microbiology, 44(5), 126247. 10.1016/j.syapm.2021.126247

Chen S, Yao H, Han J et al., (2010). Validation of the ITS2 Region as a Novel DNA Barcode for Identifying Medicinal Plant Species. Public Library of Science, 5(1), e8613. 10.1371/journal.pone.0008613

Conti I, Medrzycki P, Argenti C et al., (2016). Sugar and protein content in different monofloral pollens-building a database. Bulletin of Insectology, 69(2), 318–320.

Cook SM, Awmack CS, Murray DA et al., (2003). Are honey bees’ foraging preferences affected by pollen amino acid composition? Ecological Entomology, 28(5), 622–627. 10.1046/j.1365-2311.2003.00548.x

Crailsheim K. (1992). The Flow of Jelly within a Honeybee Colony. Journal of Comparative Physiology B, 162, 681–689. 10.1007/BF00301617

Crall JD, Chang JJ, Oppenheimer RL et al., (2017). Foraging in an unsteady world: Bumblebee flight performance in field realistic turbulence. Interface Focus, 7, 20160086. 10.1098/rsfs.2016.0086

Dahl, Å. (2018). Pollen lipids can play a role in allergic airway inflammation. Frontiers in Immunology, 9, 2816. 10.3389/fimmu.2018.02816

Debnam S, Sáez A, Aizen MA et al., (2021). Exotic insect pollinators and native pollination systems. Plant Ecology, 222, 1075–1088. 10.1007/s11258-021-01162-0

Dharampal PS, Carlson C, Currie CR et al., (2019). Pollen-borne microbes shape bee fitness. Proceedings of the Royal Society B: Biological Sciences, 286(1904), 20182894. 10.1098/rspb.2018.2894

Di Pasquale G, Salignon M, Le Conte Y et al., (2013). Influence of Pollen Nutrition on Honey Bee Health: Do Pollen Quality and Diversity Matter? Public Library of Science, 8(8), e72016. 10.1371/journal.pone.0072016

Dolezal AG and Toth AL. (2018). Feedbacks between nutrition and disease in honey bee health. Current Opinion in Insect Science, 26, 114 – 119. 10.1016/j.cois.2018.02.006

Donkersley P, Rhodes G, Pickup RW et al., (2018). Bacterial communities associated with honeybee food stores are correlated with land use. Ecology and Evolution, 8(10), 4743–4756. 10.1002/ece3.3999

Dosch C, Manigk A, Streicher T et al., (2021). The gut microbiota can provide viral tolerance in the honey bee. Microorganisms, 9(4), 871. 10.3390/microorganisms9040871

Douglas AE. (2018). Fundamentals of Microbiome Science: How Microbes Shape Animal Biology. Princeton University Press. Retrieved from: https://www.perlego.com/book/740163/fundamentals-of-microbiome-science-how-microbes-shape-animal-biology-pdf

du Rand EE, Stutzer C, Human H et al., (2020). Antibiotic treatment impairs protein digestion in the honeybee, *Apis mellifera*. Apidologie, 51, 94–106. 10.1007/s13592-019-00718-4

Encinas-Viso F, Bovill J, Albrecht DE et al., (2022). Pollen DNA metabarcoding reveals cryptic diversity and high spatial turnover in alpine plant-pollinator networks. Molecular Ecology, 32(23), 6377–6393. 10.1111/mec.16682

Engel P, Kwong WK, McFrederick Q et al., (2016). The Bee Microbiome: Impact on Bee Health and Model for Evolution and Ecology of Host-microbe Interactions. mBio, 7(2), e02164–15. 10.1128/mBio.02164-15.

Engel P, Martinson VG & Moran NA. (2012). Functional diversity within the simple gut microbiota of the honey bee. Proceedings of the Natural Academy of Sciences, 109(27), 11002– 11007. 10.1073/pnas.1202970109

Engel P and Moran NA. (2013). The gut microbiota of insects - diversity in structure and function. FEMS Microbiology Reviews, 37(5), 699–735. 10.1111/1574-6976.12025

Escalas A, Auguet JC, Avouac A, et al., (2022). Shift and homogenization of gut microbiome during invasion in marine fishes. Animal Microbiome. 4(37). 10.1186/s42523-022-00181-0

Estaki M, Jiang L, Bokulich NA et al., (2020). QIIME 2 enables comprehensive endCtoCend analysis of diverse microbiome data and comparative studies with publicly available data. Current protocols in bioinformatics, 70(1), e100. 10.1002/cpbi.100

Feás X, Vázquez-Tato MP, Estevinho L et al., (2012). Organic bee pollen: Botanical origin, nutritional value, bioactive compounds, antioxidant activity and microbiological quality. Molecules, 17(7), 8359–77. 10.3390/molecules17078359

Fick SE, and Hijmans RJ. (2017). WorldClim 2: new 1-km spatial resolution climate surfaces for global land areas. International Journal of Climatology, 37(12), 4302–4315. 10.1002/joc.5086

Figueroa LL, Maccaro JJ, Krichilsky E et al., (2021). Why did the Bee eat the chicken? Symbiont Gain, loss, and Retention in the Vulture Bee Microbiome. mBio, 12(6), e0231721. 10.1128/mBio.02317-21

Fijen TPM. (2021). Mass-migrating bumblebees: an overlooked phenomenon with potential far-reaching implications for bumblebee conservation. Journal of Applied Ecology, 58(2), 274–280. 10.1111/1365-2664.13768

Fontaine SS and Kohl KD. (2020). Gut microbiota of invasive bullfrog tadpoles responds more rapidly to temperature than a noninvasive congener. Molecular Ecology, 29(13), 2449–2462. 10.1111/mec.15487

Frankie GW, Thorp RW, Schindler M et al., (2005). Ecological patterns of bees and their host ornamental flowers in two northern California cities. Journal of the Kansas Entomological Society, 78(3), 227–246. 10.2317/0407.08.1

Garbuzov M, Alton K and Ratnieks FLW. (2017). Most ornamental plants on sale in garden centres are unattractive to flower-visiting insects. PeerJ – the Journal of Life and Environmental Sciences, 5, e3066. 10.7717/peerj.3066

Garbuzov M, Samuelson EEW and Ratnieks FLW. (2015). Survey of insect visitation of ornamental flowers in Southover grange garden, Lewes, UK. Insect Science, 22(5), 700–5. 10.1111/1744-7917.12162

Ghisbain G, Gérard M, Wood TJ et al., (2021). Expanding insect pollinators in the Anthropocene. Biological Reviews, 96(6), 2755–2770. 10.1111/brv.12777

Giurfa M, Dafni A and Neal PR. (1999). Floral symmetry and its role in plant-pollinator systems. International Journal of Plant Sciences, 160(S6). 10.1086/314214

Goulson D. (2003). Effects of introduced bees on native ecosystems. Annual Review of Ecology, Evolution and Systematics, 34, 1–26. 10.1146/annurev.ecolsys.34.011802.132355

Goulson D, Nicholls E, Botias C et al., (2015). Bee declines driven by combined stress from parasites, pesticides, and lack of flowers. Science, 347(6229), 1255957. 10.1126/science.1255957

Gray A, Brodschneider R, Adjlane N et al., (2019). Loss rates of honey bee colonies during winter 2017/18 in 36 countries participating in the COLOSS survey, including effects of forage sources, Journal of Apicultural Research, 58(4), 479–485. 10.1080/00218839.2019.1615661

Gu Y, Han W, Wang Y et al., (2023). *Xylocopa caerulea* and *Xylocopa auripennis* harbor a homologous gut microbiome related to that of eusocial bees. Frontiers in Microbiology, 14, 1124964. https://doi/10.3389/fmicb.2023.1124964

Hanley ME, Franco M, Pichon S et al., (2008). Breeding system, pollinator choice and variation in pollen quality in British herbaceous plants. Functional Ecology, 22, 592–598. 10.1111/j.1365-2435.2008.01415.x

Hanley ME and Goulson D. (2003). Introduced weeds pollinated by introduced bees: Cause or effect? Weed biology and Management, 3(4), 204–212. 10.1046/j.1444-6162.2003.00108.x

Haydak MH. (1970). Honey bee nutrition. Annual Review of Entomology, 15, 143–156. 10.1146/annurev.en.15.010170.001043

Herbert J, Elton W and Shimanuki H. (1978). Chemical composition and nutritive value of bee-collected and bee-stored pollen. Apidologie, 9(1), 33–40. 10.1051/apido:19780103

Herrera CM and Pellmyr O. (2002). Plant-animal interactions: an evolutionary approach. Blackwell Science, Oxford.

Hill K, Santosa A and England M. (2009). Interannual Tasmanian rainfall variability associated with large-scale climate modes. Journal of Climate, 22(16), 4383–4397. 10.1175/2009JCLI2769.1

Hingston AB. (2006a). Is the exotic bumblebee *Bombus terrestris* really invading Tasmanian native vegetation? Journal of insect conservation, 10(3), 289–293. 10.1007/s10841-006-6711-7

Hingston AB. (2006b). Is the introduced Bumblebee (*Bombus terrestris*) assisting the naturalization of the *Agapanthus praecox* ssp. *orientalis* in Tasmania? Ecological Management and Restoration, 7(3), 236–240. 10.1111/j.1442-8903.2006.312_7.x

Hingston AB. (2007). The potential impact of the large earth Bumblebee *‘Bombus terrestris’* (Apidae) on the Australian Mainland: lessons from Tasmania. Victorian Naturalist, 124(2), 110– 117.

Hingston AB, Marsden-Smedley JON, Driscoll DA et al., (2002). Extent of invasion of Tasmanian native vegetation by the exotic bumblebee *Bombus terrestris* (Apoidea: Apidea). Austral Ecology, 27(2), 162–172. 10.1046/j.1442-9993.2002.01179.x

Hingston AB and McQuillan PB. (1998). Nectar robbing in *Epacris impressa* (Epacridaceae) by the recently introduced bumblebee *Bombus terrestris* (Apidae) in Tasmania. Victorian Naturalist, 115, 116–119.

Hingston AB and McQuillan PB. (1999). Displacement of Tasmanian native megachilid bees by the recently introduced bumblebee Bombus terrestris (Linnaeus, 1758) (Hymenoptera: Apidae). Australian Journal of Zoology, 47(1), 59–65. 10.1071/ZO98016

Hingston AB and Wotherspoon S. (2017). Introduced social bees reduce nectar availability during the breeding season of the swift parrot (*Lathamus discolor*). Pacific Conservation Biology, 23(1), 52–62. 10.1071/PC16025

Hornick T, Richter A, Harpole WS et al., (2021). An integrative environmental pollen diversity assessment and its importance for the sustainable development goals. Plants, People, Planet, 4(2), 110–121. 10.1002/ppp3.10234

Hülsmann M, von Wehrden H, Klein A et al., (2015). Plant diversity and composition compensate for negative effects of urbanization on foraging bumble bees. Apidologie, 46, 760– 770. 10.1007/s13592-015-0366-x

Invernizzi C, Santos E, García E et al., (2011). Sanitary and nutritional characterization of honeybee colonies in *Eucalyptus grandis* plantations. Archivos de Zootecnia, 60(232), 1303– 1314. 10.4321/S0004-05922011000400045

Iovane M, Cirillo A, Izzo LG et al., (2022). High Temperature and Humidity Affect Pollen Viability and Longevity in *Olea europaea* L. Agronomy, 12(1), 1. 10.3390/agronomy12010001

Jackson JM, Pimsler ML, Oyen KJ et al., (2018). Distance, elevation and environment as drivers of diversity and divergence in bumble bees across latitude and altitude. Molecular Ecology, 27(14), 2926–2942. 10.1111/mec.14735

Jackson JM, Pimsler ML, Oyen KJ et al., (2020). Local adaptation across a complex bioclimatic landscape in two montane bumble bee species. Molecular Ecology, 29(5), 920–939. 10.1111/mec.15376

Jędrzejewska-Szmek K and Zych M. (2013). Flower-visitor and pollen transport networks in a large city: structure and properties. Arthropod-Plant Interactions, 7, 503–516. 10.1007/s11829-013-9274-z

Kakumanu ML, Reeves AM, Anderson TD et al, (2016). Honey Bee Gut Microbiome Is Altered by In-Hive Pesticide Exposures. Frontiers in Microbiology, 7. 10.3389/fmicb.2016.01255

Kantsa A, Raguso RA, Lekkas T et al., (2019). Floral volatiles and visitors: A meta-network of associations in a natural community. Journal of Ecology, 107, 2574–2586. 10.1111/1365-2745.13197

Kardum Hjort C, Paris JR, Smith HG et al., (2023a). Selection despite low genetic diversity and high gene flow in a rapid island invasion of the bumblebee, *Bombus terrestris*. Molecular Ecology, 33(2), e17212. 10.1111/mec.17212

Kardum Hjort C, Smith HG, Allen AP et al., (2023b). Morphological Variation in Bumblebees (*Bombus terrestris*) (Hymenoptera: *Apidae*) After Three Decades of an Island Invasion. Journal of insect science, 23(1). 10.1093/jisesa/iead006

Katoh K, Misawa K, Kuma KI et al., (2002). MAFFT: a novel method for rapid multiple sequence alignment based on fast Fourier transform. Nucleic acids research, 30(14), 3059–66. 10.1093/nar/gkf436

Keller A, Danner N, Grimmer G et al., (2015). Evaluating multiplexed next-generation sequencing as a method in palynology for mixed pollen samples. Plant Biology, 17(2), 558–66. 10.1111/plb.12251

Keller A, McFrederick QS, Dharampal P et al., (2021). (More than) Hitchhikers through the network: the shared microbiome of bees and flowers. Current Opinion in Insect Science, 44, 8–15. 10.1016/j.cois.2020.09.007

Knapp S, Dinsmore L, Fissore C et al., (2012). Phylogenetic and functional characteristics of household yard floras and their changes along an urbanization gradient. Ecology, 93(sp8), S83– S98. 10.1890/11-0392.1

Koch H and Schmid-Hempel P. (2011). Socially transmitted gut microbiota protect bumble bees against an intestinal parasite. Proceedings of the National Academy of Sciences, 108(48), 19288–92. 10.1073/pnas.1110474108

Koot EM, Morgan-Richards M and Trewick SA (2022). Climate change and alpine-adapted insects: Modelling environmental envelopes of a grasshopper radiation. Royal Society Open Science, 9, 211596. 10.1098/rsos.211596

Kostryco M and Chwil M. (2021). Structure of Anther Epidermis and Endothecium, Production of Pollen, and Content of Selected Nutrients from six Rubus idaeus L. Cultivars. Agronomy, 11(9), 1723. 10.3390/agronomy11091723

Krams R, Gudra D, Popovs S et al., (2022). Dominance of Fructose-Associated *Fructobacillus* in the Gut Microbiome of Bumblebees (*Bombus terrestris*) Inhabiting Natural Forest Meadows. Insects, 13(1). 10.3390/insects13010098

Kulhanek K, Steinhauer N, Rennich K et al., (2017). A national survey of managed honey bee 2015-2016 annual colony losses in the USA. Journal of Apicultural Ressearch, 56(4). 328–340. 10.1080/00218839.2017.1344496

Kwong WK, Engel P, Koch H et al., (2014). Genomics and host specialization of honey bee and bumble bee gut symbionts. Proceedings of the National Academy of Sciences, 111(31), 11509–14. 10.1073/pnas.1405838111

Kwong WK, Mancenido AL and Moran NA. (2017). Immune system stimulation by the native gut microbiota of honey bees. Royal Society Open Science, 4(2), 170003. 10.1098/rsos.170003

Kwong WK and Moran NA. (2016). Gut microbial communities of social bees. Nature Reviews Microbiology, 14, 374–384. 10.1038/nrmicro.2016.43

Lane P, Morris D, Bridle K et al., (2015). Common grasses of Tasmania. Cradle Coast NRM, NMR North, NRM South and the University of Tasmania, Hobart. 1–147. Retrieved from: https://nrmsouth.org.au/wp-content/uploads/2014/11/CommonGrassesofTasmaniaLaneetal2015.pdf

Lawson DA and Rands SA. (2019). The effects of rainfall on plant–pollinator interactions. Arthropod-Plant Interactions, 13, 561–569. 10.1007/s11829-019-09686-z

Lee FJ, Miller KI, McKinlay JB et al., (2018). Differential carbohydrate utilization and organic acid production by honey bee symbionts. FEMS Microbiology Ecology, 94(8). 10.1093/femsec/fiy113

Lee FJ, Rusch DB, Stewart FJ et al., (2015). Saccharide breakdown and fermentation by the honey bee gut microbiome. Environmental Microbiology, 17(3), 796–815. 10.1111/1462-2920.12526

Liolios V, Tananaki C, Dimou M et al., (2016). Ranking pollen from bee plants according to their protein contribution to honey bees. Journal of Apicultural Research, 54(5), 582–592. 10.1080/00218839.2016.1173353

Liu H, Hall MA, Brettell LE et al., (2023). Microbial diversity in stingless bee gut is linked to host wing size and influenced by the environment. Journal of Invertebrate Pathology, 198, 107909. 10.1016/j.jip.2023.107909

Loo WT, García-Loor J, Dudaniec RY et al., (2019). Host phylogeny, diet, and habitat differentiate the gut microbiomes of Darwin’s finches. Scientific Reports, 9, 18781. 10.1038/s41598-019-54869-6

Loram AL, Warren P, Thompson K et al., (2011). Urban domestic gardens: the effects of human interventions on garden composition. Environmental Management, 48, 808–824. 10.1007/s00267-011-9723-3

Lowenstein DM, Matteson KC and Minor ES. (2019). Evaluating the dependence of urban pollinators on ornamental, non-native, and ‘weedy’ floral resources. Urban Ecosystems, 22(1). 10.1007/s11252-018-0817-z

Lowenstein DM and Minor ES. (2016). Diversity in flowering plants and their characteristics: integrating humans as a driver of urban floral resources. Urban Ecosystems, 19, 1735–1748. 10.1007/s11252-016-0563-z

Lozupone C, Stombaugh J, Gordon J et al., (2012). Diversity, stability and resilience of the human gut microbiota. Nature, 489, 220–230. 10.1038/nature11550

Lucas A, Bodger O, Brosi BJ et al., (2018). Generalisation and specialisation in hoverfly (Syrphidae) grassland pollen transport networks revealed by DNA metabarcoding. Journal of Animal Ecology, 87(4), 1008–1021. 10.1111/1365-2656.12828

Macfarlane RP and Gurr L. (1995). Distribution of bumble bees in New Zealand. New Zealand Entomologist, 18(1), 29–36. 10.1080/00779962.1995.9721999

Manfredini F, Arbetman M and Toth AL, (2019). A potential role for phenotypic plasticity in invasions and declines in social insects. Fronters in Ecology and Evolution, 7. 10.3389/fevo.2019.00375

Manning R. (2001). Fatty acids in pollen: A review of their importance for honey bees. Bee World, 82(2), 60–75. 10.1080/0005772X.2001.11099504

Martinez Arbizu, P. (2020). pairwiseAdonis: Pairwise multilevel comparison using adonis. R package, version 0.4. Retrieved from: https://github.com/pmartinezarbizu/pairwiseAdonis

Martin M. (2011). Cutadapt removes adapter sequences from high-throughput sequencing reads. EMBnet.journal, 17(1), 10–12, 10.14806/ej.17.1.200

Martinson VG, Danforth BN, Minckley RL et al., (2011). A simple and distinctive microbiota associated with honey bees and bumble bees. Molecular Ecology, 20(3), 619–28. 10.1111/j.1365-294X.2010.04959.x

Matteson KC and Langellotto GA. (2009). Bumble bee abundance in New York City community gardens: implications for urban agriculture. Cities and the Environment, 2(1), 5, Retrieved from: https://digitalcommons.lmu.edu/cate/vol2/iss1/5

McFrederick QS and Rehan SM. (2022). Wild Bee Pollen Usage and Microbial Communities Co-vary Across Landscapes. Microbial Ecology, 77(2), 513–522. 10.1007/s00248-018-1232-y

McFrederick QS, Thomas JM, Neff JL et al., (2017). Flowers and Wild Megachilid Bees Share Microbes. Microbial Ecology, 73(1), 188–200. 10.1007/s00248-016-0838-1

McMurdie PJ and Holmes S. (2013). phyloseq: an R package for reproducible interactive analysis and graphics of microbiome census data. Public Library of Science, 8(4), e61217. 10.1371/journal.pone.0061217

Milla, L, SchmidtCLebuhn A, Bovill, J et al., (2022). Monitoring of honey bee floral resources with pollen DNA metabarcoding as a complementary tool to vegetation surveys. Ecological solutions and evidence, 3(1), e12120. 10.1002/2668-8319.12120

Motta EVS and Moran NA. (2024). The honeybee microbiota and its impact on health and disease. Nature Reviews Microbiology, 22(3), 122–137. 10.1038/s41579-023-00990-3

Mountcastle AM, Ravi S and Combes SA. (2015). Nectar vs. pollen loading affects the tradeoff between flight stability and maneuverability in bumblebees. Proceedings of the National Academy of Sciences, 112(33), 10527–10532. 10.1073/pnas.1506126112

Naug D. (2009). Nutritional stress due to habitat loss may explain recent honeybee colony collapses. Biological Conservation, 142(10), 2369–2372. 10.1016/j.biocon.2009.04.007

Neumann P and Carreck NL. (2010). Honey bee colony losses. Journal of Apicultural Research, 49(1), 1–6. 10.3896/IBRA.1.49.1.01

Nicolson SW (2011). Bee Food: The Chemistry and Nutritional Value of Nectar, Pollen and Mixtures of the Two. African Zoology, 46, 197–204. 10.3377/004.046.0201

O’Connell M, Jordan Z, McGilvray E et al., (2021). Reap what you sow: local plant composition mediates bumblebee foraging patterns within urban garden landscapes. Urban Ecosystems, 24(5), 391–404. 10.1007/s11252-020-01043-w

Oksanen J, Blanchet FG, Friendly M et al., (2024). Package ‘vegan’. Community ecology package, version 2.6-4. Retrieved from: https://cran.r-project.org/web/packages/vegan/vegan.pdf

Pinheiro J, Bates D, DebRoy S, Sarkar D and R Core Team. (2024). nlme: Linear and Nonlinear Mixed Effects Models, *version 3.1-166*. Retrieved from: https://cran.r-project.org/web/packages/nlme/nlme.pdf

Pornon A, Andalo C, Burrus M et al., (2017). DNA metabarcoding data unveils invisible pollination networks. Scientific Reports, 7, 16828. 10.1038/s41598-017-16785-5

Potts SG, Biesmeijer JC, Kremen C et al., (2010). Global pollinator declines: trends, impacts and drivers. Trends in Ecology and Evolution, 25(6), 345–353. 10.1016/j.tree.2010.01.007

Price MN, Dehal PS and Arkin AP. (2010). FastTree 2 – Approximately Maximum-Likelihood Trees for Large Alignments. Public Library of Science, 5(3), e9490. 10.1371/journal.pone.0009490

Quast C, Pruesse E, Yilmaz P et al., (2012). The SILVA ribosomal RNA gene database project: improved data processing and web-based tools. Nucleic acids research, 41 (Database issue), D590–D596. 10.1093/nar/gks1219

R Core Team. (2024). R: A language and Environment for Statistical Computing. R Foundation for Statistical Computing, Vienna, Austria [Methodology Reference]. Retrieved from: https://www.R-project.org/

Requier F, Antúnez K, Morales CL et al., (2018). Trends in beekeeping and honey bee colony losses in Latin America. Journal of Apicultural Research, 57(5), 1–6. 10.1080/00218839.2018.1494919

Ricigliano VA, Fitz W, Copeland DC et al., (2017). The impact of pollen consumption on honey bee (*Apis mellifera*) digestive physiology and carbohydrate metabolism. Archives of Insect Biochemistry and Physiology, 96(2). 10.1002/arch.21406

Ricigliano VA, Williams ST and Oliver R. (2022). Effects of different artificial diets on commercial honey bee colony performance, health biomarkers, and gut microbiota. BMC Veterinary Research, 18, 52. 10.1186/s12917-022-03151-5

Robeson MS, O’Rourke DR, Kaehler BD et al., (2021). RESCRIPt: Reproducible sequence taxonomy reference database management. Public Library of Science: Computational Biology, 17(11), e1009581. 10.1371/journal.pcbi.1009581

Roulston TH and Buchmann SL. (2000). A phylogenetic reconsideration of the pollen starch-pollination correlation. Evolutionary Ecology Research, 2(5), 627–643. Retrieved from: https://www.evolutionary-ecology.com/issues/v02n05/ggar1186.pdf

Roulston TH and Cane JH. (2000). Pollen nutritional content and digestibility for animals. Plant Systematics and Evolution, 222, 187–209. 10.1007/BF00984102

Rudra Gouda MN, Deeksha MG, Kumarang KM et al., (2024). Diversity of Gut Microbes in the Forager and Hive Bees of an Indian Population of *Apis mellifera*. National Academy Science Letters, 47(2). 10.1007/s40009-023-01318-8

Russo L. (2016). Positive and negative impacts of non-native bee species around the world. Insects, 7(4), 69. 10.3390/insects7040069

Saa-Otero MP; Diaz-Losada E and Fernandez-Gomez E (2000). Analysis of fatty acids, proteins and ethereal extract in honeybee pollen-considerations of their floral origin. Grana, 39(4), 175–181. 10.1080/00173130051084287

Sáez A, Morales CL, Garibaldi LA et al., (2017). Invasive bumble bees reduce nectar availability for honey bees by robbing raspberry flower buds. Basic and Applied Ecology, 19, 26–35. 10.1016/j.baae.2017.01.001

Scarth P. (2013). Vegetation Height and Structure - Derived from ALOS-1 PALSAR, Landsat and ICESat/GLAS, Australia Coverage. Terrestrial Ecosystem Research Network (TERN). Dataset. Retrieved from: https://portal.tern.org.au/metadata/TERN/de1c2fef-b129-485e-9042-8b22ee616e66

Schmid-Hempel P, Schmid-Hempel R, Brunner PC et al., (2007). Invasion success of the bumblebee, *Bombus terrestris*, despite a drastic genetic bottleneck. Heredity, 99(4), 414–22. 10.1038/sj.hdy.6801017

Schwarz RS, Moran NA, and Evans JD. (2016). Early gut colonizers shape parasite susceptibility and microbiota composition in honey bee workers. Proceedings of the National Academy of Sciences, 113(33), 9345–50. 10.1073/pnas.1606631113

Semmens TD, Turner E and Buttermore R. (1993). *Bombus terrestris* (L.) (Hymenoptera: Apidae) now established in Tasmania. Australian Journal of Entomology, 32(4). 10.1111/j.1440-6055.1993.tb00598.x

Siepielski AM, Morrissey MB, Buoro M et al., (2017). Precipitation drives global variation in natural selection. Science, 355(6328), 959–962. 10.1126/science.aag2773

Singh T, Bhat MM and Khan MA. (2009). Insect adaptations to changing environments – Temperature and humidity. International Journal of Industrial Entomology, 19(1), 155–164. Retrieved from: https://koreascience.kr/article/JAKO200933063804084.pdf

Somerville DC. (2001). Nutritional value of bee collected pollens. Rural Industries Research and Development Corporation, 1–166. Retrieved from: https://www.nbba.ca/wp-content/uploads/2013/12/Nutritional_Value_of_Bee_Collected_Pollens.pdf

Stanimirović Z, Glavinić U, Ristanić M et al., (2019). Looking for the causes of and solutions to the issue of honey bee colony losses. Acta Veterinaria, 69(1), 1–31. 10.2478/acve-2019-0001

Steffan SA and Dharampal PS. (2019). Undead foodCwebs: Integrating microbes into the foodCchain. Food Webs, 18, e00111. 10.1016/j.fooweb.2018.e00111

Steffan SA, Dharampal PS, Danforth BN et al., (2019). Omnivory in bees: Elevated trophic positions among all major bee families. American Naturalist, 194, 3, 414–421. 10.1086/704281

Steinhauer N, Kulhanek K, Antúnez K et al., (2018). Drivers of colony losses. Current Opinion in Insect Science, 26, 142–148. 10.1016/j.cois.2018.02.004

Tang Q-H, Li W-L, Wang J-P et al., (2023). Effects of spinetoram and glyphosate on physiological biomarkers and gut microbes in *Bombus terrestris*. Fronters in Physiology, 13. 10.3389/fphys.2022.1054742

Tsadila C, Amoroso C and Mossialos D. (2023). Microbial Diversity in Bee Species and Bee Products: Pseudomonads Contribution to Bee Well-Being and the Biological Activity Exerted by Honey Bee Products: A Narrative Review. Diversity, 15(10), 1088. 10.3390/d15101088

Vanderplanck M, Leroy B, Wathelet B et al., (2014). Standardized protocol to evaluate pollen polypeptides as bee food source. Apidologie, 45(2), 192–204. 10.1007/s13592-013-0239-0

Vannette RL, Gauthier MPL and Fukami T. (2012). Nectar bacteria, but not yeast, weaken a plantCpollinator mutualism. Proceedings of the Royal Society B: Biological Sciences, 280, 20122601. 10.1098/rspb.2012.2601

Vásquez A, Forsgren E, Fries I et al., (2012). Symbionts as major modulators of insect health: lactic acid bacteria and honeybees. Public Library of Science, 7(3), e33188. 10.1371/journal.pone.0033188

Winston ML. (1991). The Biology of the Honey Bee. Harvard University Press, Cambridge, Massachusetts. Retrieved from: https://www.perlego.com/book/1133854/the-biology-of-the-honey-bee-pdf

Wright GA, Nicolson SW and Shafir S. (2018). Nutritional physiology and ecology of honey bees. Annual Review of Entomology. 63, 327–344. 10.1146/annurev-ento-020117-043423

Yao H, Song J, Liu C et al., (2010). Use of ITS2 region as the universal DNA barcode for plants and animals. Public Library of Science, 5(10), e13102. 10.1371/journal.pone.0013102

Yilmaz P, Parfrey LW, Yarza P et al., (2014). The SILVA and ‘all-species living tree project (LTP)’ taxonomic frameworks. Nucleic acids research, 42 (Database issue), D643–8. 10.1093/nar/gkt1209

Zanette LRS, Martins RP and Ribeiro SP. (2005). Effects of urbanization on Neotropical wasp and bee assemblages in a Brazilian metropolis. Landscape and Urban Planning, 71(2-4), 105–121. 10.1016/j.landurbplan.2004.02.003

Zhang Z, Mu X, Cao Q et al., (2022). Honeybee gut *Lactobacillus* modulates host learning and memory behaviors via regulating tryptophan metabolism. Nature Communications, 13, 2037. 10.1038/s41467-022-29760-0

Zheng H, Perreau J, Powell JE et al., (2019). Division of labor in honey bee gut microbiota for plant polysaccharide digestion. Proceedings of the National Academy of Sciences, 116(51), 25909–25916. 10.1073/pnas.1916224116

Zheng H, Powell JE, Steele MI et al., (2017). Honeybee gut microbiota promotes host weight gain via bacterial metabolism and hormonal signaling. Proceedings of the National Academy of Sciences, 114(18), 4775–4780. 10.1073/pnas.1701819114

Zhu L, Zhang Z, Chen H, et al., (2021). Gut microbiomes of bigheaded carps and hybrids provide insights into invasion: A hologenome perspective. Evolutionary Applications, 14(3), 735–745. 10.1111/eva.13152.

