## Supplementary material for "Landscape-wide metabarcoding of the invasive bumblebee *(Bombus terrestris)* shows interactions among the gut microbiome and pollenbiome": Text S1

**Text S1. DNA metabarcoding library preparation and sequencing**

16S V4 region amplicon sequencing was undertaken by the Ramaciotti Centre for Genomics (University of New South Wales, Sydney, Australia). The gene-specific full-length primer sequences, targeting the V4 region were:

16S Amplicon PCR Forward Primer = 5’ TCGTCGGCAGCGTCAGATGTGTATAAGAGACAGCCTACGGGNGGCWGCAG and 16S Amplicon PCR Reverse Primer = 5’ GTCTCGTGGGCTCGGAGATGTGTATAAGAGACAGGACTACHVGGGTATCTAATCC. The Illumina overhang adapter sequences added to the locus‐specific primer for the target region: Forward overhang: 5’ TCGTCGGCAGCGTCAGATGTGTATAAGAGACAG‐[locus specific sequence] and Reverse overhang: 5’GTCTCGTGGGCTCGGAGATGTGTATAAGAGACAG‐[locus specific sequence].

To amplify the 16S gene, PCR conditions comprised an initial denaturation step at 95°C for 3 minutes, followed by 25 cycles of 95°C for 30 seconds, 55°C annealing for 30 seconds, 72°C extension for 30 seconds, and a final extension at 72°C for 5 minutes. Following this PCR, purification of the PCR products was executed using AMPure XP beads to eliminate free primers and primer dimers from the amplicons. Subsequently, a dual indexing process and attachment of Illumina sequencing adapters were conducted through an index PCR, utilizing the Nextera XT Index Kit (FC-131-1001). The conditions for this PCR began with 95°C for 3 minutes which was followed by 8 cycles of 95°C for 30 seconds, 55°C annealing for 30 seconds, 72°C for 30 seconds, and a final extension at 72°C for 5 minutes. A secondary PCR clean-up step utilizing AMPure XP beads was performed before the library underwent final quantification and validation. A 1:50 dilution of the ultimate library was ran on a Bioanalyzer DNA 1000 chip to confirm the size. The pooled libraries were subsequently denatured with NaOH, diluted with hybridization buffer, and heat-denatured before paired-end 2x250 sequencing on the Illumina MiSeq platform.

**SUPPLEMENTARY TABLES**

**Table S1.** Pearson’s correlation matrix for all environmental variables.

Mean seasonal precipitation was excluded from further analyses due to its notable strong positive correlation (r > 0.7) with mean annual precipitation (r = 0.71). The final six selected environmental variables are emphasized in bold. Abbreviations: Annual Temp = Mean annual temperature ($℃)$, Annual Precip = Mean annual precipitation (mm), Seasonal Precip = Precipitation seasonality (mm), Percent Pasture = Percentage of pasture (%), Veg Height = Height of vegetation (mm), Percent Urban = Percentage of urbanization (%) and Wind = Average velocity of summer wind (m/s).

|  | ***Annual Temp*** | ***Annual Precip*** | Seasonal Precip | ***Percent Pasture*** | ***Veg Height*** | ***Percent Urban*** | ***Wind*** |
| --- | --- | --- | --- | --- | --- | --- | --- |
| ***Annual Temp*** | 1 | -0.01 | 0.37 | -0.06 | -0.05 | 0.28 | 0.63 |
| ***Annual Precip*** | -0.01 | 1 | *0.71* | -0.41 | 0.47 | -0.19 | 0.23 |
| Seasonal Precip | 0.37 | 0.71 | 1 | -0.35 | 0.29 | -0.04 | 0.31 |
| ***Percent Pasture*** | -0.06 | -0.41 | -0.35 | 1 | -0.51 | -0.2 | -0.13 |
| ***Veg Height*** | -0.05 | 0.47 | 0.29 | -0.51 | 1 | 0.04 | 0.46 |
| ***Percent Urban*** | 0.28 | -0.19 | -0.04 | -0.2 | 0.04 | 1 | 0.25 |
| ***Wind*** | 0.63 | 0.23 | 0.31 | -0.13 | 0.46 | 0.25 | 1 |

**Table S2.** Linear mixed-effect model and interactions between *B. terrestris* gut bacterial richness, pollen packet richness and environmental variables. Abbreviations: Bacterial_richness = Chao1 richness estimate of gut bacterial samples, Pollen_richness = Chao1 richness estimate of pollenbiome samples, AT = Mean annual temperature (℃), AR = Mean annual precipitation (mm), PP = Percentage of pasture (%), VH = Height of vegetation (mm), PU = Percentage of urbanization (%), and WV = Average velocity of summer wind (m/s).

| Random effect = ~1\|Sites | | | |
| --- | --- | --- | --- |
| Fixed effect(s) | DF | t-value | p-value |
| Bacterial_richness~AT | 12 | 0.20 | 0.85 |
| Bacterial_richness~AR | 12 | 0.99 | 0.34 |
| Bacterial_richness~PP | 12 | -1.07 | 0.30 |
| Bacterial_richness~PU | 12 | 0.55 | 0.60 |
| Bacterial_richness~VH | 12 | -0.31 | 0.76 |
| Bacterial_richness~WV | 12 | -1.18 | 0.26 |
| Bacterial_richness~AT*AR | 10 | -0.56 | 0.59 |
| Bacterial_richness~AT*PP | 10 | -0.28 | 0.79 |
| Bacterial_richness~AT*PU | 10 | 0.93 | 0.37 |
| Bacterial_richness~AT*VH | 10 | -1.29 | 0.22 |
| Bacterial_richness~AT*WV | 10 | -0.02 | 0.98 |
| Bacterial_richness~AR*PP | 10 | -0.92 | 0.38 |
| Bacterial_richness~AR*PU | 10 | 0.20 | 0.84 |
| Bacterial_richness~AR*VH | 10 | 1.07 | 0.31 |
| Bacterial_richness~AR*WV | 10 | -0.36 | 0.73 |
| Bacterial_richness~PP*PU | 10 | -0.89 | 0.39 |
| Bacterial_richness~PP*VH | 10 | -1.03 | 0.32 |
| Bacterial_richness~PP*WV | 10 | -0.06 | 0.95 |
| Bacterial_richness~PU*VH | 10 | -0.84 | 0.42 |
| Bacterial_richness~PU*WV | 10 | -0.54 | 0.60 |
| Bacterial_richness~VH*WV | 10 | -1.18 | 0.26 |
| Bacterial_richness~Pollen_richness | 12 | 0.16 | 0.87 |
| Bacterial_richness~Pollen_richness*AT | 10 | 0.13 | 0.90 |
| Bacterial_richness~Pollen_richness*AR | 10 | -1.39 | 0.19 |
| Bacterial_richness~Pollen_richness*PP | 10 | -0.11 | 0.91 |
| Bacterial_richness~Pollen_richness*PU | 10 | -1.32 | 0.21 |
| Bacterial_richness~Pollen_richness*VH | 10 | 0.91 | 0.38 |
| Bacterial_richness~Pollen_richness*WV | 10 | 0.86 | 0.41 |

**Table S3.** Summary of pairwise PERMANOVA conducted to analyse significance of *B. terrestris* gut bacterial community composition between sites. Cells with p-values emphasized in bold indicates statistically significant sites (p < 0.05).

|  | S1 | S2 | S4 | S5 | S6 | S9 | S15 | S17 | S18 | S19 | S20 | S22 | S23 | S24 | S25 | S26 |
| --- | --- | --- | --- | --- | --- | --- | --- | --- | --- | --- | --- | --- | --- | --- | --- | --- |
| S1 |  |  |  |  |  |  |  |  |  |  |  |  |  |  |  |  |
| S2 | 0.862 |  |  |  |  |  |  |  |  |  |  |  |  |  |  |  |
| S4 | 0.177 | 0.222 |  |  |  |  |  |  |  |  |  |  |  |  |  |  |
| S5 | **0.021** | **0.006** | 0.241 |  |  |  |  |  |  |  |  |  |  |  |  |  |
| S6 | 0.165 | 0.117 | 0.37 | 0.147 |  |  |  |  |  |  |  |  |  |  |  |  |
| S9 | **0.012** | **0.007** | **0.019** | 0.6 | **0.035** |  |  |  |  |  |  |  |  |  |  |  |
| S15 | 0.7 | 0.69 | 0.236 | **0.044** | 0.191 | **0.012** |  |  |  |  |  |  |  |  |  |  |
| S17 | 0.09 | 0.068 | 0.051 | 0.067 | **0.026** | **0.005** | 0.653 |  |  |  |  |  |  |  |  |  |
| S18 | 0.174 | 0.246 | 0.812 | 0.191 | 0.403 | **0.003** | 0.467 | 0.052 |  |  |  |  |  |  |  |  |
| S19 | 0.065 | 0.176 | 0.501 | 0.05 | 0.124 | **0.001** | 0.102 | **0.021** | 0.544 |  |  |  |  |  |  |  |
| S20 | **0.007** | **0.002** | 0.053 | 0.079 | **0.027** | **0.005** | **0.013** | **0.007** | 0.2 | 0.298 |  |  |  |  |  |  |
| S22 | 0.054 | 0.06 | **0.024** | **0.004** | **0.008** | 0.194 | **0.016** | **0.004** | **0.018** | **0.005** | **0.004** |  |  |  |  |  |
| S23 | 0.14 | **0.048** | 0.292 | 0.058 | 0.074 | **0.002** | 0.142 | **0.047** | 0.277 | 0.136 | 0.097 | **0.005** |  |  |  |  |
| S24 | 0.064 | 0.055 | **0.029** | **0.031** | **0.033** | **0.016** | 0.644 | 0.973 | **0.023** | **0.022** | **0.006** | **0.008** | **0.012** |  |  |  |
| S25 | 0.574 | 0.734 | 0.702 | **0.023** | 0.626 | **0.012** | 0.768 | 0.096 | 0.674 | 0.16 | **0.007** | **0.019** | 0.098 | 0.076 |  |  |
| S26 | 0.245 | 0.131 | 0.144 | 0.086 | 0.609 | **0.04** | 0.4 | 0.218 | 0.115 | **0.045** | **0.009** | **0.006** | **0.02** | 0.2 | 0.552 |  |

**Table S4.** Summary of *t-test* conducted to analyse the significance of differences in Chao1 richness index of *B. terrestris* gut bacteria between sites. All sites are statistically insignificant (p > 0.05).

|  | S1 | S15 | S17 | S18 | S19 | S2 | S20 | S22 | S23 | S24 | S25 | S26 | S4 | S5 | S6 |
| --- | --- | --- | --- | --- | --- | --- | --- | --- | --- | --- | --- | --- | --- | --- | --- |
| S15 | 1 |  |  |  |  |  |  |  |  |  |  |  |  |  |  |
| S17 | 1 | 1 |  |  |  |  |  |  |  |  |  |  |  |  |  |
| S18 | 1 | 1 | 1 |  |  |  |  |  |  |  |  |  |  |  |  |
| S19 | 1 | 1 | 1 | 1 |  |  |  |  |  |  |  |  |  |  |  |
| S2 | 1 | 1 | 1 | 1 | 1 |  |  |  |  |  |  |  |  |  |  |
| S20 | 1 | 1 | 1 | 1 | 1 | 1 |  |  |  |  |  |  |  |  |  |
| S22 | 1 | 1 | 1 | 1 | 1 | 1 | 1 |  |  |  |  |  |  |  |  |
| S23 | 1 | 1 | 1 | 1 | 1 | 0.083 | 0.913 | 0.979 |  |  |  |  |  |  |  |
| S24 | 1 | 1 | 1 | 1 | 1 | 1 | 1 | 1 | 1 |  |  |  |  |  |  |
| S25 | 1 | 1 | 1 | 1 | 1 | 0.684 | 1 | 1 | 1 | 1 |  |  |  |  |  |
| S26 | 1 | 1 | 1 | 1 | 1 | 1 | 1 | 1 | 1 | 1 | 1 |  |  |  |  |
| S4 | 1 | 1 | 1 | 1 | 1 | 1 | 1 | 1 | 0.182 | 1 | 1 | 1 |  |  |  |
| S5 | 1 | 1 | 1 | 1 | 1 | 1 | 1 | 1 | 0.351 | 1 | 1 | 1 | 1 |  |  |
| S6 | 1 | 1 | 1 | 1 | 1 | 1 | 1 | 1 | 1 | 1 | 1 | 1 | 1 | 1 |  |
| S9 | 1 | 1 | 1 | 1 | 1 | 1 | 1 | 1 | 1 | 1 | 1 | 1 | 1 | 1 | 1 |

**Table S5.** Summary of ANOVA conducted to analyse the significance of differences in Shannon’s diversity index of *B. terrestris* gut bacteria between sites. Cells with p-values emphasized in bold shows statistically significant sites (p < 0.05).

|  | S1 | S2 | S4 | S5 | S6 | S9 | S15 | S17 | S18 | S19 | S20 | S22 | S23 | S24 | S25 |
| --- | --- | --- | --- | --- | --- | --- | --- | --- | --- | --- | --- | --- | --- | --- | --- |
| S1 |  |  |  |  |  |  |  |  |  |  |  |  |  |  |  |
| S2 | 0.999 |  |  |  |  |  |  |  |  |  |  |  |  |  |  |
| S4 | 1 | 1 |  |  |  |  |  |  |  |  |  |  |  |  |  |
| S5 | 0.999 | 0.996 | 0.999 |  |  |  |  |  |  |  |  |  |  |  |  |
| S6 | 0.999 | 0.958 | 0.999 | 1 |  |  |  |  |  |  |  |  |  |  |  |
| S9 | 0.308 | 0.087 | 0.349 | 0.723 | 0.913 |  |  |  |  |  |  |  |  |  |  |
| S15 | 1 | 1 | 1 | 0.999 | 0.997 | 0.283 |  |  |  |  |  |  |  |  |  |
| S17 | 0.997 | 0.894 | 0.994 | 0.999 | 1 | 0.987 | 0.986 |  |  |  |  |  |  |  |  |
| S18 | 0.999 | 1 | 1 | 0.994 | 0.947 | 0.078 | 1 | 0.876 |  |  |  |  |  |  |  |
| S19 | 0.998 | 1 | 0.999 | 0.898 | 0.688 | **0.01** | 0.999 | 0.554 | 1 |  |  |  |  |  |  |
| S20 | 0.999 | 1 | 0.999 | 0.974 | 0.865 | **0.032** | 1 | 0.754 | 1 | 1 |  |  |  |  |  |
| S22 | 0.999 | 0.988 | 0.999 | 1 | 1 | 0.736 | 0.999 | 0.999 | 0.983 | 0.814 | 0.941 |  |  |  |  |
| S23 | 0.999 | 0.996 | 0.999 | 1 | 1 | 0.624 | 0.999 | 0.999 | 0.994 | 0.895 | 0.975 | 1 |  |  |  |
| S24 | 0.9 | 0.532 | 0.888 | 0.997 | 0.999 | 0.999 | 0.835 | 1 | 0.499 | 0.162 | 0.322 | 0.999 | 0.994 |  |  |
| S25 | 0.999 | 0.966 | 0.999 | 1 | 1 | 0.895 | 0.998 | 1 | 0.957 | 0.72 | 0.885 | 1 | 1 | 0.999 |  |
| S26 | 0.763 | 0.357 | 0.762 | 0.982 | 0.999 | 0.999 | 0.689 | 0.999 | 0.329 | 0.082 | 0.187 | 0.986 | 0.964 | 1 | 0.998 |

**Table S6.** List of plant types foraged by *B. terrestris* across each Tasmanian site. Explanation of type of plant categories: ‘Native genera’ = plant genera native to Tasmania, ‘Introduced genera’ = plant genera that are exotic and have been naturalized in Tasmania, ‘Both genera’ = plant genera consisting of both native and introduced species in Tasmania.

| **Introduced genera** | **Both genera** | **Native genera** |
| --- | --- | --- |
| Rubus | Plantago | Eucalyptus |
| Hypochaeris | Solanum | Myrcia |
| Lupinus | Veronica | Leptospermum |
| Lotus | Borago | Clematis |
| Raphanus | Melaleuca | Correa |
| Reseda | Sonchus | Myoporum |
| Sisymbrium | Epilobium |  |
| Pisum | Senecio |  |
| Trifolium | Acacia |  |
| Rosa | Erigeron |  |
| Digitalis |  |  |
| Osmanthus |  |  |
| Prunus |  |  |
| Brassica |  |  |
| Cirsium |  |  |
| Glycine Max |  |  |
| Circaea |  |  |
| Cichorium |  |  |
| Syzigium |  |  |
| Medicago |  |  |
| Sanguisorba |  |  |
| Cosmos |  |  |
| Centaurium |  |  |
| Vigna |  |  |
| Tropaeolum |  |  |
| Achillea |  |  |
| Anagalis |  |  |
| Kickxia |  |  |
| Leucanthemum |  |  |
| Arnoseris |  |  |
| Buddleja |  |  |
| Hydrangea |  |  |
| Reynoutria |  |  |
| Wisteria |  |  |
| Corymbia |  |  |

**SUPPLEMENTARY FIGURES**

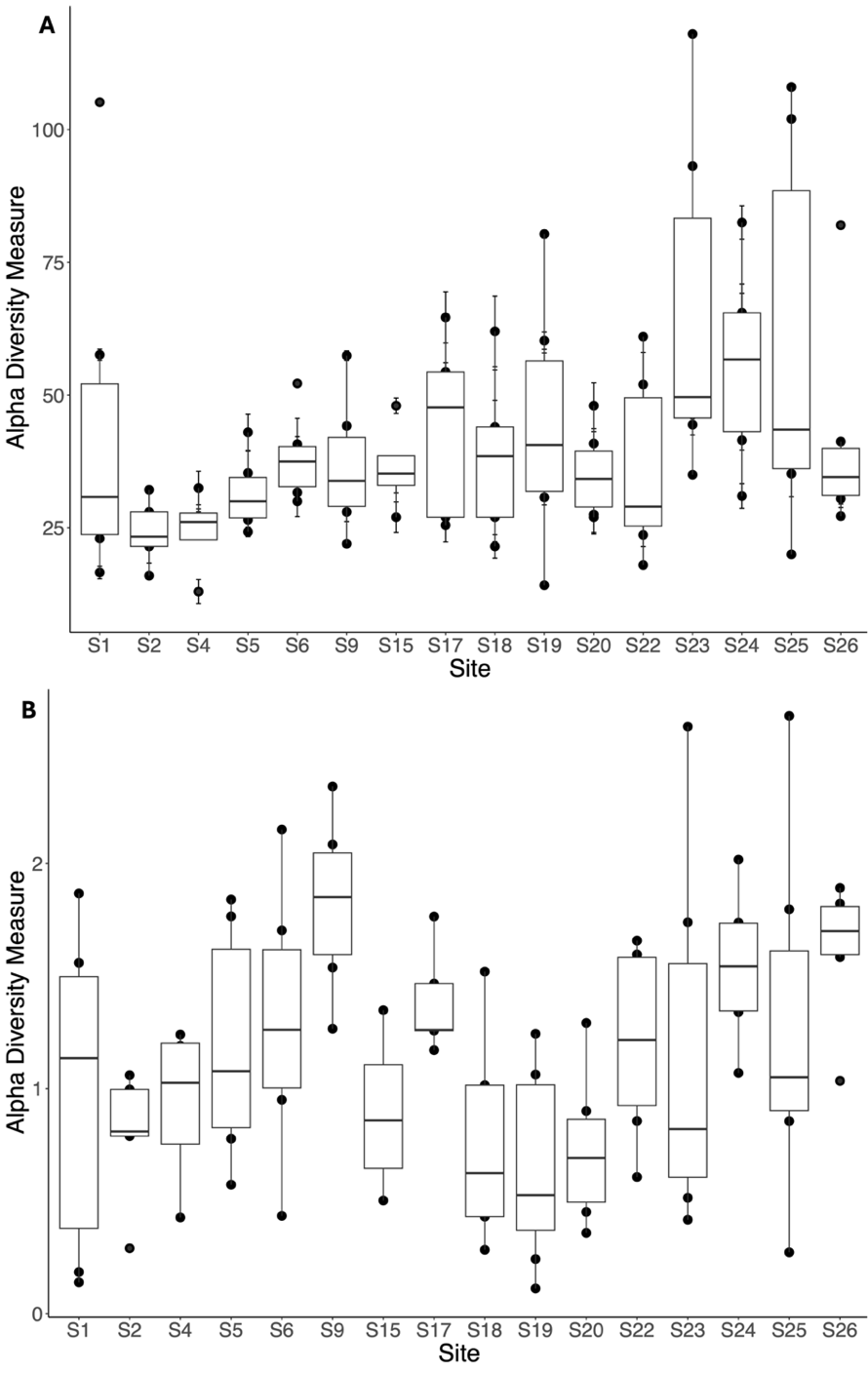

**Fig S1.** Alpha diversity measures for *B. terrestris* gut bacteria across 16 sites. (A) Chao1 richness estimates for *B. terrestris* gut bacterial samples per site. All sites were statistically insignificant, *t-test*: p > 0.05 (see Table S3 for all corresponding t-test results) (B) Shannon’s diversity indices for *B. terrestris* gut bacterial samples per site. S9 is statistically significant with S19, ANOVA: p = 0.01; S9 is statistically significant with S20, ANOVA: p = 0.032 (see Table S4 for all corresponding ANOVA results). For both plots, boxes represent the median, standard deviation and outliers.

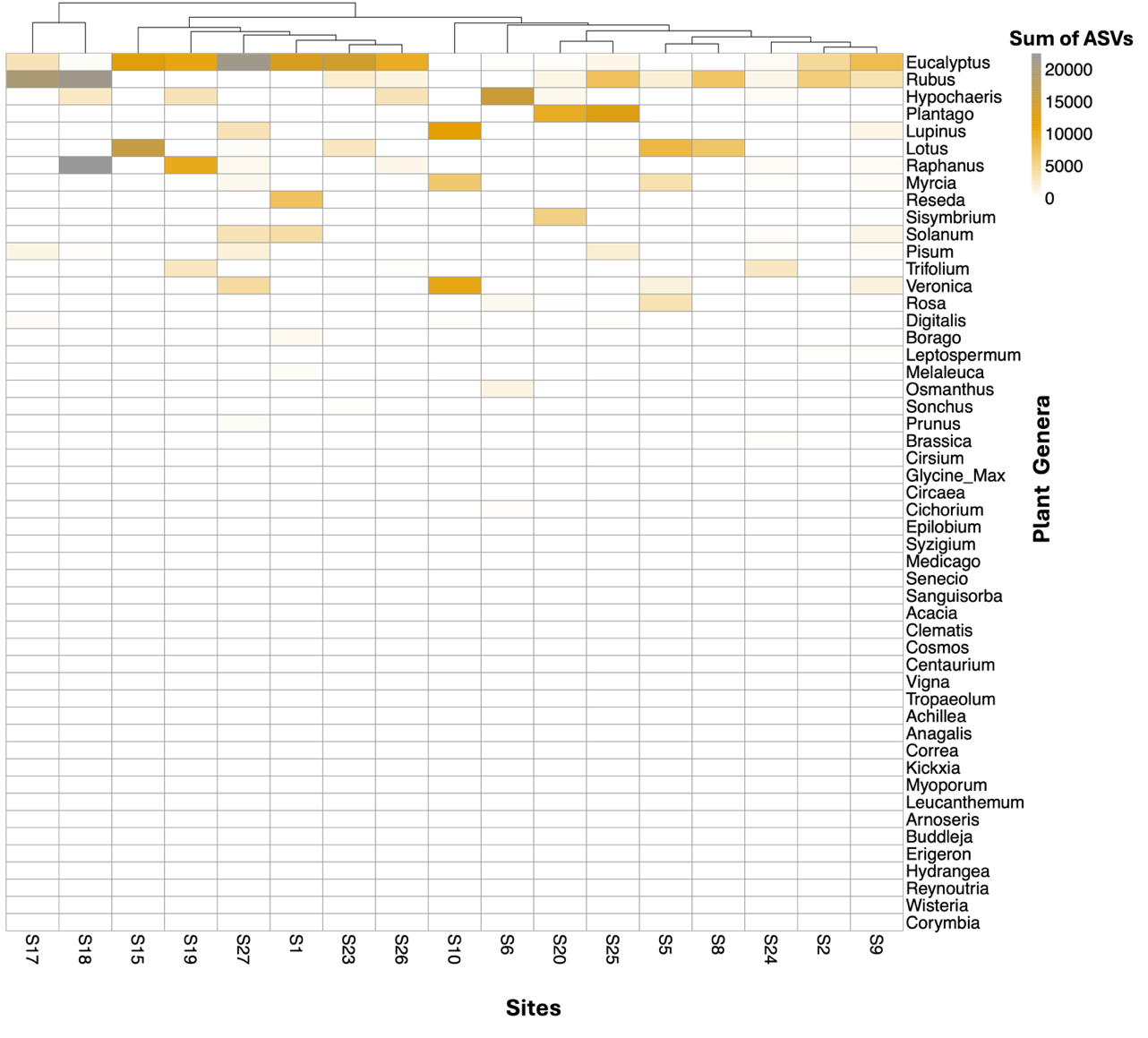

**Fig S2.** Heatmap showing all the plant genera identified from the pollen packets of *B. terrestris* across 17 sites. The colour scales indicate the sum of amplicon sequence variants (ASVs) under each plant genus. The dendrogram on the top of the heatmap shows the distance or similarities among the sites. The dendrogram is generated using Euclidean distance as the hierarchial clustering measure, which revealed the specific nodes for each site.

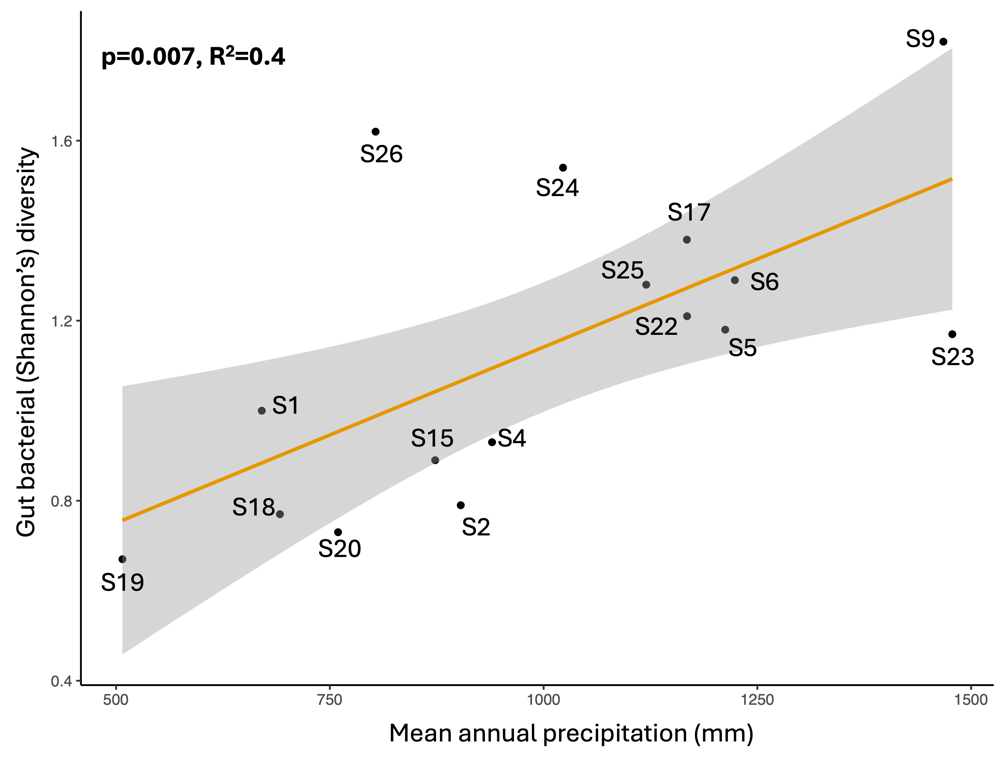

**Fig S3.** The positive effect of mean annual precipitation on gut bacterial diversity of *B. terrestris* across Tasmanian sites.

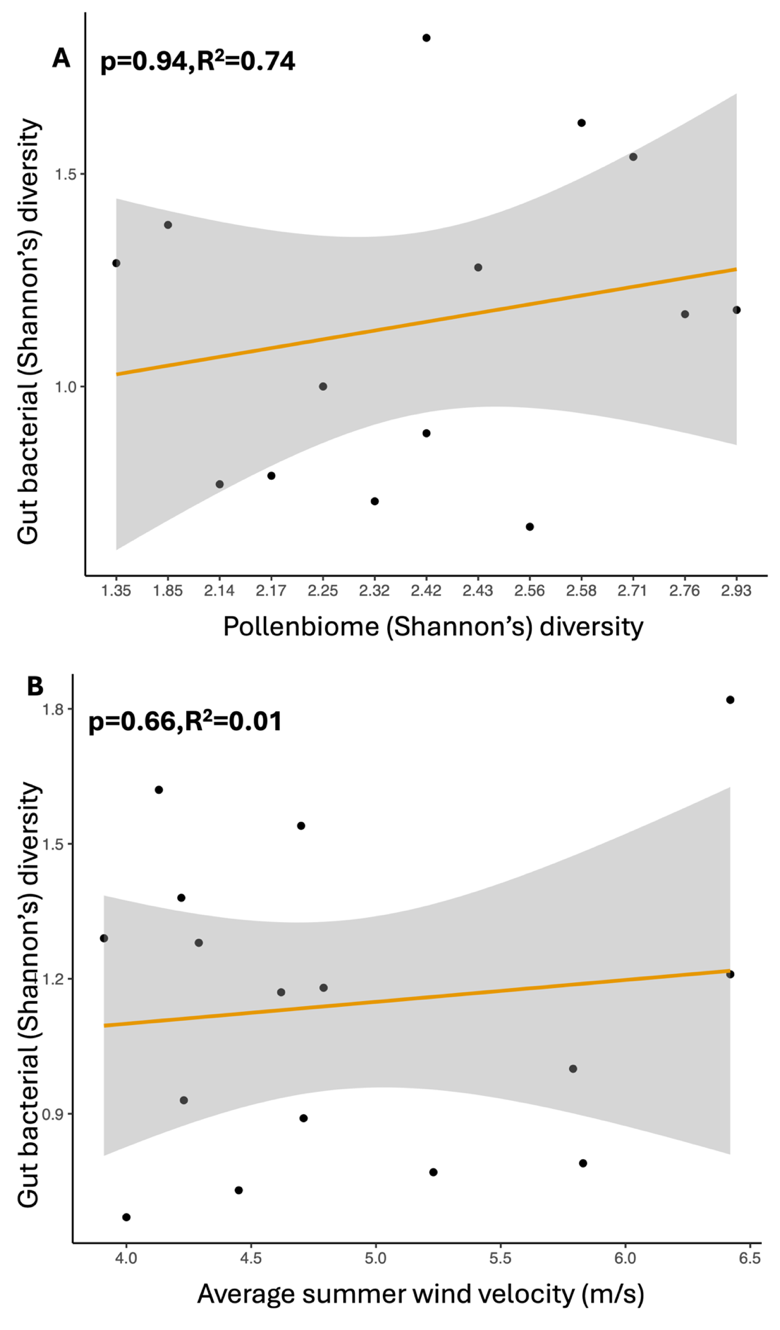

**Fig S4.** (A) Relationship between gut bacterial diversity and pollen packet diversity of *B. terrestris*. (B) Relationship between *B. terrestris* gut bacterial diversity and average summer wind velocity. Both linear relationships are statistically insignificant (*lm*: p > 0.05). In both plots, the dots represent different Tasmanian sites.

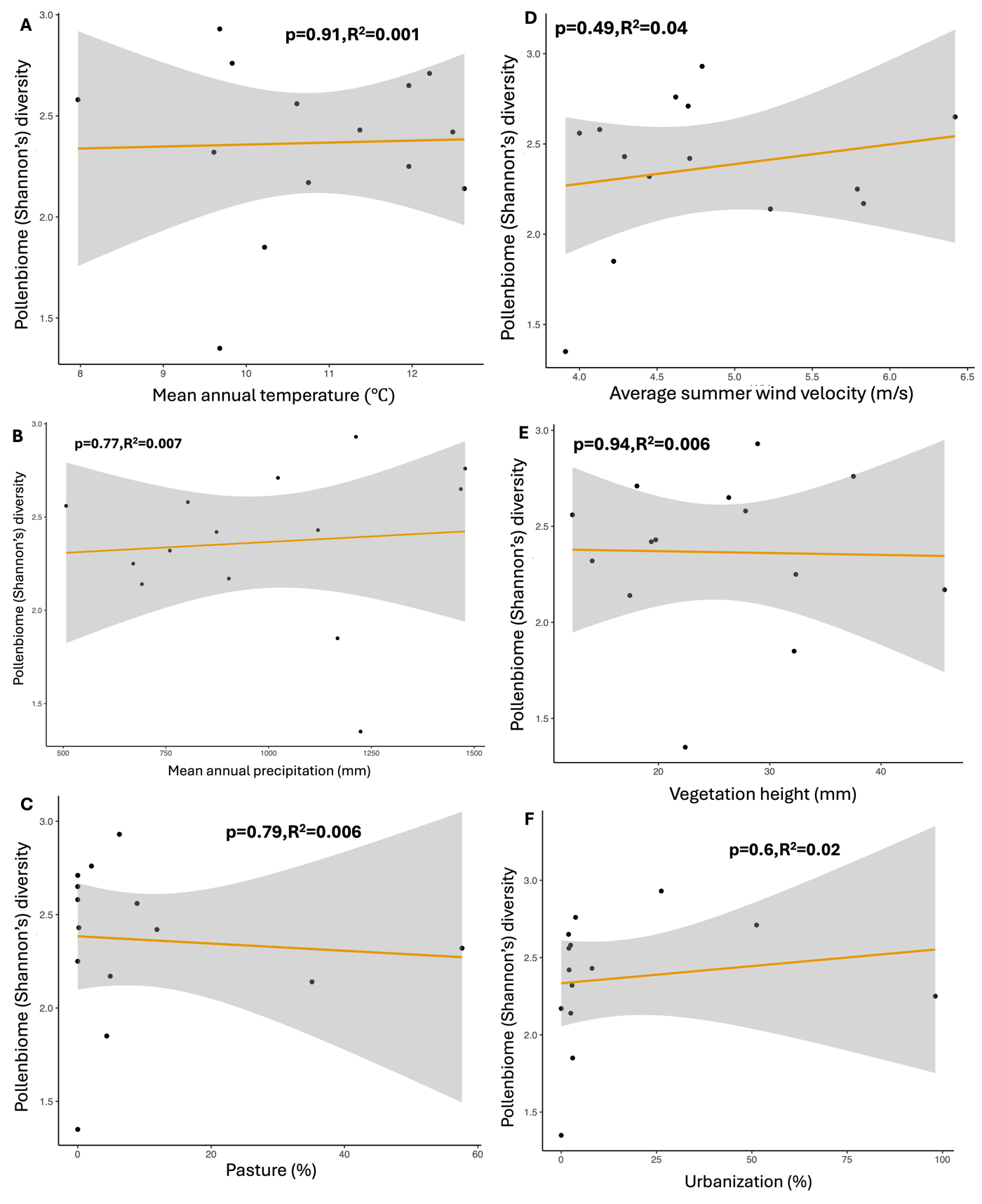

**Fig S5.** Relationship between Shannon’s diversity of pollen packets and the six environmental variables which includes: (A) mean annual temperature, (B) mean annual precipitation, (C) percentage of pasture, (D) average summer wind velocity, (E) height of vegetation and (F) percentage of urbanization. All six linear plots indicate statistically insignificant interactions between pollen diversity and environmental factors (*lm*: p > 0.05). In both plots, the dots represent different Tasmanian sites.

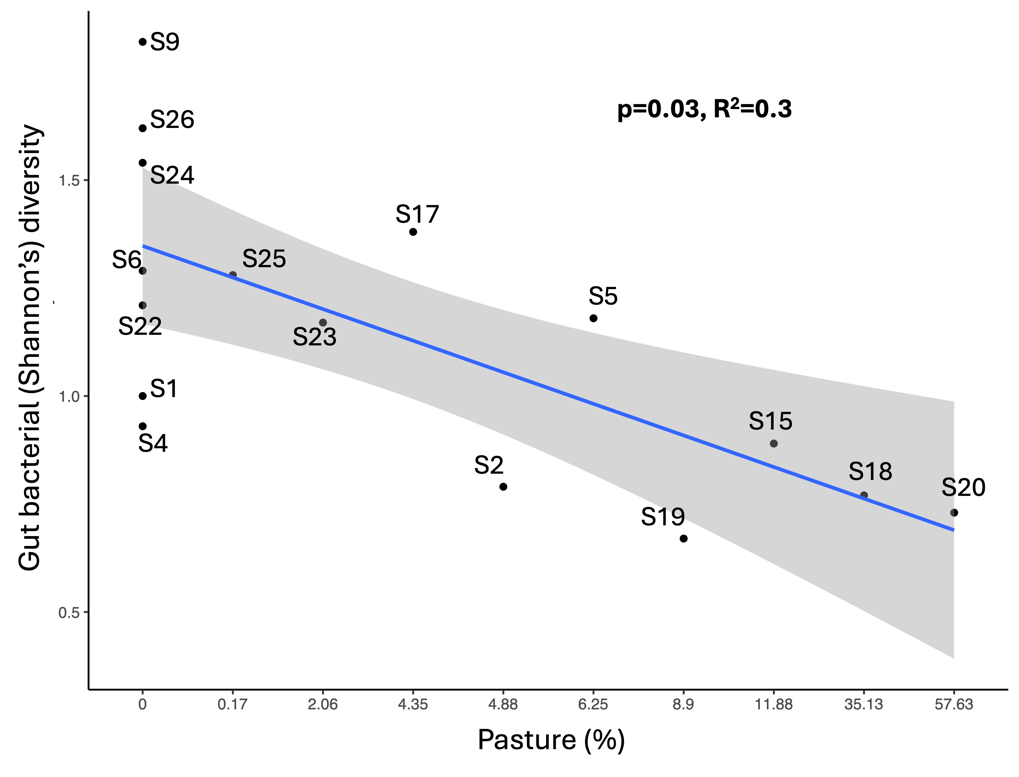

**Fig S6.** The negative effect of percentage of pasture on diversity of *B. terrestris* gut bacteria across Tasmanian sites.
